# Land use intensification favours fitness similarity as plant diversity declines

**DOI:** 10.1101/2022.12.15.520291

**Authors:** Rodrigo R. Granjel, Oscar Godoy, Fons van der Plas, Santiago Soliveres, Caterina Penone, Hugo Saiz, Norbert Hölzel, Daniel Prati, Markus Fischer, Eric Allan

## Abstract

1. Land use intensification reduces plant diversity, but we know little about how it changes coexistence mechanisms. Niche and fitness differences, together with indirect interactions, can respond non-linearly to intensification and influence which species combinations are feasible, meaning that all members have positive equilibrium abundances.
2. We analysed nine annual vegetation surveys (2008–2016) of 50 common plant species across 150 German grasslands, using a composite land use intensity (LUI) index summarising fertilisation, mowing and grazing. We used population models and a structural approach to estimate feasibility, niche and fitness differences along the LUI gradient, and examined indirect effects in assemblages of three to eleven species. To investigate why some species were absent locally, we compared locally observed pairs and triplets with combinations containing one locally absent species. We also tested how well feasible combinations reconstructed richness and distinguished locally present from absent species, using both the fitted interactions and a scenario with facilitation removed.
3. Across the regional pool, feasibility tended to decline and fitness differences to increase with LUI, but neither change was statistically significant. Niche differences remained high, with an intermediate peak in triplets. Locally observed combinations gained advantages in feasibility and fitness similarity over unobserved alternatives as LUI increased. They also had higher niche differences, but this advantage weakened with intensification. Feasibility depended more strongly on indirect effects in larger assemblages, particularly at low and high LUI. Richness reconstructed with competition and facilitation initially declined but then rebounded, exceeding the observed maximum throughout. Removing facilitation brought estimates closer to the observed maximum and its decline with LUI.
4. *Synthesis.* Our results suggest that land use intensification favours fitness similarity among species found together locally as plant diversity declines. Time series can help reveal changes in coexistence mechanisms and guide experiments varying management intensity and resident neighbours to identify what limits diversity recovery.

## Introduction

Increases in land use intensity (LUI), through greater fertilisation, earlier or more frequent mowing, or more intensive grazing, are a major threat to biodiversity (Rockström et al., 2009). Although many studies have shown that plant diversity declines (Allan et al., 2014; Crawley et al., 2005; Newbold et al., 2015) and species composition changes (Dormann et al., 2007; Pakeman, 2011) as land use intensifies, we have a poor understanding of the mechanisms responsible for these processes, which hampers efforts to predict and manage land use effects. Numerous studies have used changes in functional diversity to investigate the mechanisms of community assembly. Some have found that communities at high land use intensity become dominated by functionally similar species (Laliberté & Tylianakis, 2012; Mayfield et al., 2010; Pakeman, 2011). For example, the narrowing ranges of canopy height, leaf dry matter content and seed mass associated with land use intensification have been interpreted as environmental filtering (Pakeman, 2011). However, it can be challenging to infer precise coexistence mechanisms from functional trait distributions (Mayfield & Levine, 2010). An alternative is to take a demographic approach in which changes in population abundance are analysed to estimate how species interactions and environmental conditions affect whether species can persist together over time (Hallett et al., 2019; Song, Von Ahn, et al., 2020; Tredennick et al., 2017).

Changes in environmental conditions can alter species’ intrinsic growth rates and the strength of competitive and facilitative interactions (Adler, Smull, et al., 2018; HilleRisLambers et al., 2012; Olsen et al., 2016). If altered environmental conditions in intensively managed grasslands result in negative intrinsic growth rates, this may lead to extinction (Selwood et al., 2015). However, whether reduced intrinsic growth rates lead to species loss may also depend on neighbouring plants, as facilitative effects from neighbours may help to sustain populations that would otherwise go extinct (Buche et al., 2025; Losapio et al., 2021). Land use intensification may also alter competitive interactions, alongside intrinsic growth rates, and therefore affect two complementary components of coexistence (Chesson, 2000; Saavedra et al., 2017). In classical models of competition, niche differences promote coexistence by causing species to limit themselves more strongly than they limit others, whereas smaller fitness differences can favour coexistence by reducing asymmetries between species (Fig. 1). Together, intrinsic growth rates and species interactions determine whether a combination is feasible, meaning that all members have positive equilibrium abundances (Adler, Smull, et al., 2018; Saavedra et al., 2017). We use feasibility to assess opportunities for coexistence, while recognising that positive equilibrium abundances alone do not establish persistence through time. Although non-linear changes in diversity with land use intensification have been documented (Allan et al., 2014; Saiz et al., 2026), we still know little about how opportunities for coexistence change along these gradients or whether these responses are also non-linear.

**Figure 1.**
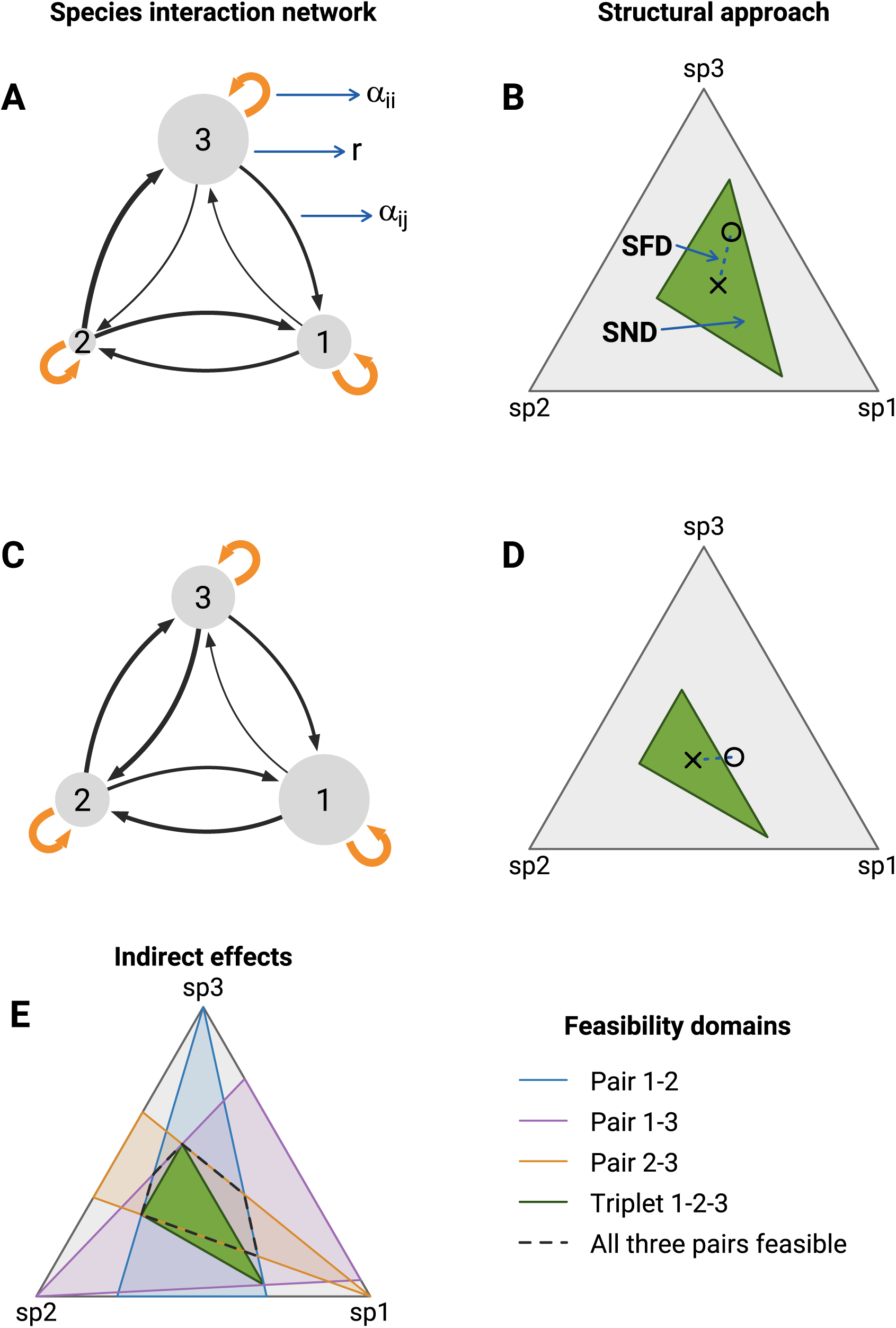
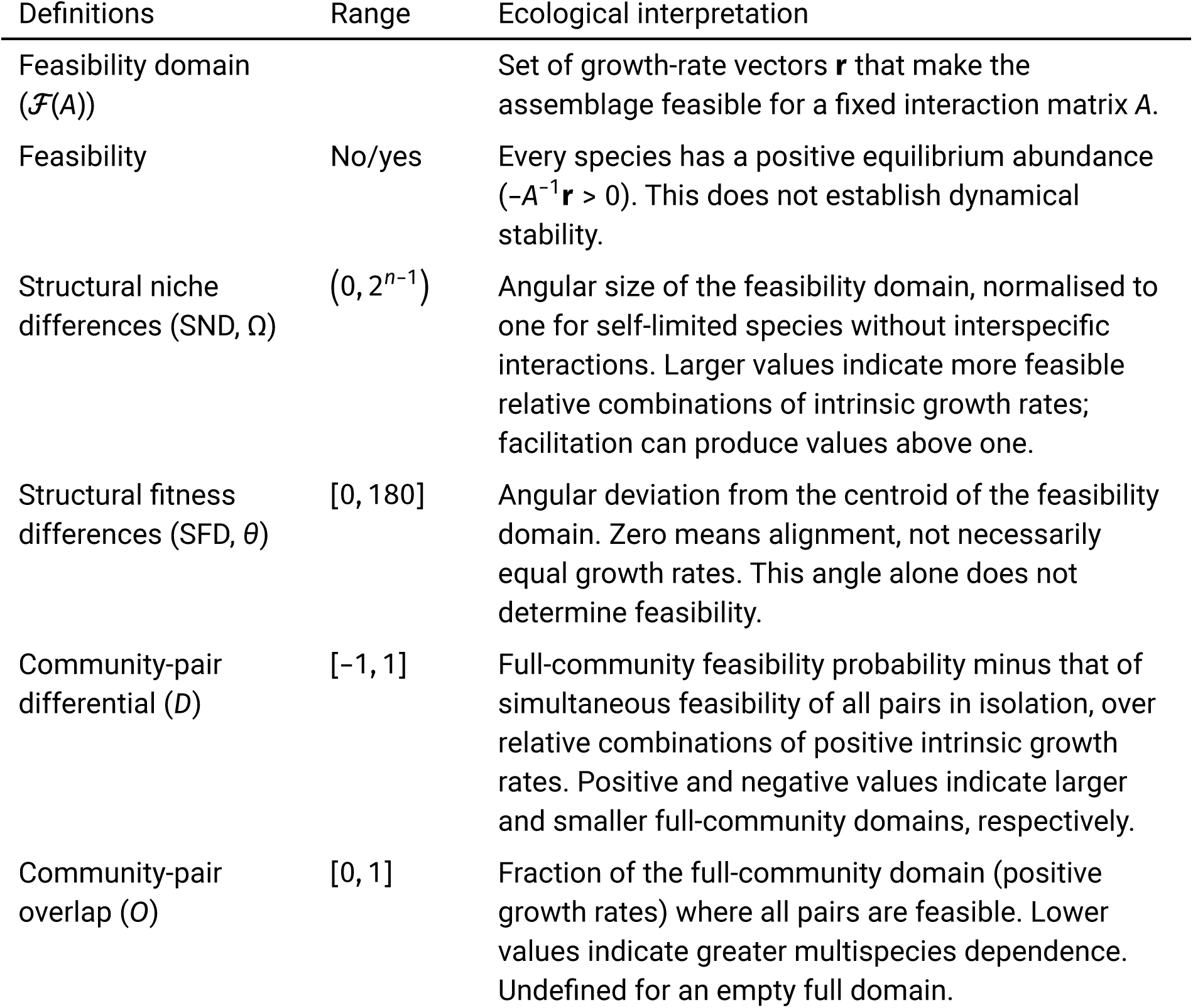
Species interactions and feasibility for two hypothetical three-species competitive assemblages. (A,C) Orange loops show intraspecific competition and dark arrows interspecific competition (*α_ij_* < 0). Thicker lines indicate stronger effects; larger circles indicate higher intrinsic growth rates (*r*). In A, the labels show species 3 acting on itself (*α_ii_* = *α*_33_) and on species 1 (*α_ij_* = *α*_13_). (B,D) Green feasibility domains are projected onto the grey positive-growth simplex. Open circles mark growth-rate vectors: feasible in B, infeasible in D. Crosses mark the centroid of the feasibility domain (**r***_c_*), and dotted blue lines schematically indicate its angular separation from the growth-rate vector (SFD). SND measures normalised solid angle, not planar area. (E) The matrix from C–D defines the pairwise domains (colours as in the legend) and triplet domain (green). Each pair’s domain leaves the absent species’ growth rate unconstrained; the dashed outline marks simultaneous feasibility of all three pairs. In this representation, the differential metric compares the spherical sizes of the green and dashed domains, whereas the overlap metric is the fraction of the green domain inside the dashed domain. The table applies to invertible interaction matrices with *n* ≥ 2 species. With facilitation, feasibility domains can extend beyond the positive simplex. SND and SFD are not directly compared to determine feasibility. Equations are given in the Supplementary Material, Methods. Diagrams adapted from Petry & Lepori (2022), following Saavedra et al. (2017).

Various aspects of land use intensification can alter species’ niche and fitness differences. For example, fertilisation can increase asymmetry in competition for light, as taller plants intercept a disproportionate share of light and shade smaller neighbours (DeMalach et al., 2017). This could increase fitness differences between species that benefit from nutrient enrichment and those that become increasingly shaded. Fertilisation can also change the balance between below-ground and above-ground competition by relieving nutrient limitation (Wilson & Tilman, 1993), which could reduce niche differences if species coexist through differences in nutrient use. At the same time, mowing and grazing may increase light levels for low-growing species and therefore reduce differences in competitive ability (Eskelinen et al., 2022). Fertilisation, mowing and grazing could therefore have different effects on coexistence. Many changes in coexistence mechanisms are possible as land use is intensified, and a key unanswered question is how these detailed mechanisms together shape overall niche and fitness differences along the LUI gradient. If we can estimate how species interactions change with land use intensity, we can also test whether coexistence models reproduce the corresponding changes in species richness. This is a key test for coexistence models.

In multispecies communities, species can also coexist through indirect interactions, which can influence feasibility in several ways. One example is intransitive competition, e.g., the rock-paper-scissors game, in which no species is a universally superior competitor (Laird & Schamp, 2006; Soliveres et al., 2015). Beyond this clear example, weaker forms of intransitivity are possible, which could promote coexistence in combination with niche differences (Gallien et al., 2017; Godoy et al., 2017). Another example is indirect facilitation, in which species A indirectly benefits C by suppressing C’s competitor B (Godoy et al., 2017; Miller, 1994). Land use may alter the importance of these indirect interactions for feasibility (Soliveres et al., 2015). For example, fertilisation can increase competitive asymmetry (DeMalach et al., 2017; Wilson & Tilman, 1993), potentially creating more hierarchical networks in which a few species dominate the others because differences in individual size can translate into unequal competitive effects (Rees, 2013). Land use can also alter direct facilitative effects and thereby change the indirect effects transmitted through an assemblage (Mayfield et al., 2010). However, how land use intensification changes opportunities for coexistence and their dependence on indirect interactions remains unclear.

In summary, land use intensification could reduce opportunities for coexistence by increasing fitness differences, reducing niche differences or weakening the benefits of indirect interactions. To examine these mechanisms, we can use coexistence models to estimate their responses across potential species combinations in a regional pool. However, patterns across the whole pool may differ from those among species actually found together locally. The species remaining at high land use intensity might still coexist, for instance through high niche differences, so analysing only these species could miss changes associated with species loss. Theoretical and empirical work has shown that observed combinations are more likely to persist under environmental change than many potential alternatives from the same regional species pool (Medeiros et al., 2021). We can therefore ask whether the species that remain as management intensifies form combinations with greater feasibility, stronger niche differentiation and greater fitness similarity than alternative combinations from the regional pool that are not observed locally. These comparisons can help us investigate possible explanations for why some species from the regional pool are absent from local communities.

Here, we address these questions using annual records of plant cover from 2008 to 2016 in 150 grasslands of the German Biodiversity Exploratories (Fischer et al., 2010). We focus on the 50 most common plant taxa and use cover as a proxy for abundance. These grasslands differ in their fertilisation, mowing and grazing intensity, summarised by a composite LUI index (Blüthgen et al., 2012). Species richness declines with increasing LUI (Fig. S1), consistent with patterns reported elsewhere (Crawley et al., 2005; Flynn et al., 2009; Newbold et al., 2015). We apply a structural stability approach (Rohr et al., 2014; Saavedra et al., 2017) that quantifies the range of intrinsic growth-rate combinations that allow an assemblage to be feasible, termed the feasibility domain (Fig. 1). A larger domain indicates more opportunities for coexistence (Godoy et al., 2018; Saavedra et al., 2017). We examine how feasibility and structural analogues of niche and fitness differences vary with LUI. We calculate these measures for pairs and triplets of species using intrinsic growth rates and interaction coefficients predicted at different levels of land use intensity, considering the combined index and each management component separately. To relate these patterns to the species observed in local communities, we compare observed combinations occurring at a given land use level with combinations from the regional pool that do not locally occur. We then assess how indirect interactions alter feasibility in larger assemblages and how strongly their effects depend on the number of interacting species. Finally, we test whether we can reconstruct observed richness by counting the distinct species represented in feasible pairs and triplets. We also assess whether species’ participation in these combinations distinguishes locally present from absent species. Both assessments compare the fitted interactions, which include competition and facilitation, with a scenario in which facilitative effects are removed. Removing the facilitation component can help determining whether observed species richness can be explained primarily by species showing positive intrinsic growth rate, which is the only mechanisms of persisting when population dynamics are governed exclusively by competition.

## Materials and Methods

### Study system and data

The Biodiversity Exploratories (www.biodiversity-exploratories.de) project (Fischer et al., 2010) is a research initiative that has established 150 permanent grassland plots of 50 × 50 m in three different regions across Germany: the UNESCO Biosphere Area Schwäbische Alb (south-west), Hainich-Dün (central) and the UNESCO Biosphere Reserve Schorfheide-Chorin (north-east). All three regions have a similar climate, with a range of 3 °C in mean annual temperature and 500 to 1,000 mm annual precipitation (see Fischer et al. (2010) for details). These 150 plots have been managed as grasslands for at least 20 years prior to the start of the project. Farmers and landowners provide information on the intensity of land management activities, including fertilisation, mowing, and grazing (Vogt et al., 2019). Other actions such as the application of pesticides occur in a non-systematic way. These sites, which range from seminatural to intensively managed grasslands, are mown or grazed. Grazed plots have cattle, horses, or sheep, with varying numbers of animals and duration of grazing. Grazing intensity was quantified as the number of livestock-unit days ha^−1^ year^−1^. Mown plots are cut one to four times per year. Some grasslands are also fertilised, the intensity of which is quantified as the amount of organic and inorganic nitrogen added to the grasslands, in kg N ha^−1^ year^−1^ (Blüthgen et al., 2012).

With this detailed land use information, prior work has quantified a compound index of land use intensity (LUI), which integrates the intensity of fertilisation (F), the mowing frequency (M), and the intensity of grazing (G) for each grassland plot (Allan et al., 2014; Blüthgen et al., 2012; Fischer et al., 2010). Higher values of LUI indicate greater overall land use intensity through greater nitrogen inputs, more frequent mowing and/or increased grazing intensity. For example, a frequently mown, fertilised meadow and an intensively grazed, fertilised pasture can have similar LUI values (quantitative details in Allan et al. (2014)). The main advantage of this index is that it summarises different correlated aspects of land use into a single metric (Table S1). Here, we used annual LUI records for 2008–2015, with values ranging between 0 and 4.57 across the plots (Fig. S1). The underlying management records are publicly available (Vogt et al., 2019).

The vegetation database used here comprised nine annual surveys from 2008 to 2016, in which species percentage cover was recorded in permanent 4 × 4 m subplots at peak biomass (Socher et al., 2012). Vegetation records are archived in the Biodiversity Exploratories Information System (BExIS) (Bolliger et al., 2021). A total of 345 plant taxa were recorded across the 150 plots. We paired each census from 2008–2015 with LUI in the same year and with cover in the following year, using the 2016 survey for the final annual change. For further analyses, we selected the 50 most common plant taxa, defined as those with the highest mean cover across the 2008–2015 censuses. Their scientific names and occurrence frequencies across regions and LUI levels are provided in Table S2.

### Estimating growth rates and species interactions

We modelled changes in relative plant cover to estimate how species’ growth rates and the effects of conspecific and heterospecific neighbours varied with LUI. We analysed eight annual changes in relative cover, from 2008–2009 through 2015–2016, matching each change to LUI in its starting year (2008–2015). Species percentage cover was converted to relative cover by dividing each species’ cover by the total cover of all recorded plant taxa in the same plot and year.

We fitted Gaussian linear mixed-effects models to estimate population parameters for the 50 taxa selected for the structural analyses (Table S2). The logarithmic ratio of relative cover between time *t* + 1 and time *t* was modelled as a linear function of LUI, together with the cover of each modelled taxon, including the focal taxon itself, at time *t*. We further included interactions between LUI and each cover predictor, which allowed the effects of neighbours to change along the management gradient (Clark et al., 2020; Tredennick et al., 2017). Related regression approaches have been used to predict changes in species abundances over time (Clark et al., 2020). Each model used 1,136 complete annual transitions across the 150 plots, covering the full observed LUI range. Models included random intercepts for region and plot nested within region, and accounted for temporal autocorrelation using a first-order autoregressive residual correlation structure within plots. Models were fitted by restricted maximum likelihood (REML) using the function `lme` in the `nlme` R package (Pinheiro et al., 2023). The general form of the statistical model was:

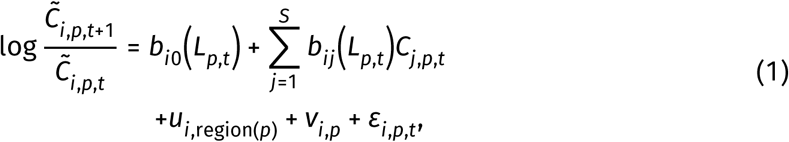

where *S* is the number of taxa included as cover predictors and *C_i_*_,*p*,*t*_ is the relative cover of taxon *i* in plot *p* and year *t*. For the logarithmic ratio, we used *C̃_i_*_,*p*,*t*_ = *C_i_*_,*p*,*t*_ + 0.01 to retain zero- cover observations. The 0.01 offset was not added to the cover predictors. *L_p_*_,*t*_ is LUI in that plot and year, and *u_i_*_,region(*p*)_ and *v_i_*_,*p*_ are the regional and plot random intercepts.

The fitted growth baseline, *b_i_*_0_(*L*), varies linearly with LUI and represents the expected log change in relative cover at zero cover of all modelled taxa, with random effects set to zero. We interpreted this baseline as a model-based estimate of intrinsic growth rate, although the lack of experimental data does not allow to corroborate it empirically. The coefficients *b_ij_*(*L*) describe the effects of conspecific and heterospecific cover on growth and also vary linearly with LUI. Positive values indicate facilitative effects and negative values competitive effects. Each focal taxon was modelled separately, so each directed cover effect could respond differently to LUI. We first summarised how intrinsic growth rates and intraspecific and interspecific effects varied with LUI and calculated the percentage of pairs and triplets containing at least one facilitative interspecific effect (Fig. S2). We used a joint parametric bootstrap to compare these summaries between LUI 0.5 and 3, with full details given in the Supplementary Material, Methods S1.

Finally, the residual term (*ε_i_*_,*p*,*t*_) accounts for variation unexplained by the fitted effects, which may include cover-measurement error (Cheng & Van Ness, 1999), dispersal and other unmodelled processes. We did not specify a term for dispersal from outside the plots because no information was available to assess propagule pressure at the plot level. These processes are therefore not separately identified by the residual structure.

### A structural approach to modelling species feasibility

We assessed how species feasibility varied along the LUI gradient using a discrete Lotka- Volterra population model (Saavedra et al., 2017):

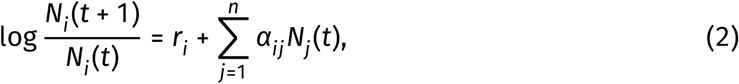

where *n* is the number of species in an assemblage, *N_i_* and *N_j_* are their relative covers, *r_i_* is the intrinsic growth rate of species *i*, and *α_ij_* is the effect of species *j* on the growth of species *i*.

Given that there is a direct correspondence between the structure of the statistical model (Equation 1) and that of the population model (Equation 2), we parameterised the population model using the fitted coefficients from the statistical model, setting *r_i_*(*L*) = *b_i_*_0_(*L*) and *α_ij_*(*L*) = *b_ij_*(*L*). We evaluated the same fitted growth-rate and interaction functions at eleven LUI values from 0.5 to 3.0, in increments of 0.25, and calculated feasibility and structural niche and fitness differences at each LUI value.

The structural approach has been applied to understand how species within ecological communities persist despite variable population dynamics (Allen-Perkins et al., 2023; Bartomeus et al., 2021; Granjel et al., 2023; Granjel et al., 2026; Song, Rohr, et al., 2020; Tabi et al., 2020; Wang et al., 2024). It allows us to examine how competitive and facilitative interactions combine to determine feasibility, and extends from pairs of species to multispecies assemblages, where indirect effects can arise. Building on the distinction between niche and fitness differences (Chesson, 2000; Godoy & Levine, 2014), the structural approach provides geometric analogues that describe how species interactions and intrinsic growth rates relate to feasibility (Saavedra et al., 2017).

We evaluated feasibility by requiring all equilibrium abundances to be positive, **N**^*^ = −*A*^−1^**r** > 0, where *A* = (*α_ij_*) is the interaction matrix and **r** the vector of intrinsic growth rates. The set of growth-rate vectors that meets this condition is the feasibility domain (Rohr et al., 2014). Feasibility is necessary, but does not by itself establish stable coexistence (Grilli et al., 2017; Saavedra et al., 2017). Structural niche differences (SND) measure the normalised angular size of this domain. Larger values indicate that a broader range of relative combinations of intrinsic growth rates is compatible with feasibility. Structural fitness differences (SFD) measure the angular deviation of the fitted growth-rate vector from the centroid of the feasibility domain. Larger values indicate greater deviation from this centroid. We tested feasibility directly, rather than by comparing SND with SFD, because their direct comparison is not a general feasibility criterion for networks that include facilitative interactions. Fig. 1 illustrates the approach, and equations and symbol definitions are provided in the Supplementary Material, Methods S2.

We calculated feasibility, niche differences and fitness differences for all 1,225 pairs and 19,600 triplets drawn from the pool of 50 taxa. The same species and combinations were evaluated at the 11 LUI values considered. We summarised feasibility as the percentage of combinations with positive equilibrium abundances. Niche and fitness differences were summarised with empirical medians, interquartile ranges and decile distributions. We interpreted these two metrics separately for pairs and triplets because their geometries change with assemblage size (Granjel et al., 2023).

Pairs and triplets share species and fitted coefficients, so they do not provide independent observations for testing changes along the LUI gradient. We therefore used a joint parametric bootstrap to account for uncertainty in the fitted population parameters when making these comparisons. We simulated new growth responses from the fitted population models, then refitted the models and recalculated the metrics, as described in the Supplementary Material, Methods S1.1. We compared feasibility between LUI 0.5 and 3 and median niche and fitness differences among LUI 0.5, 1.75 and 3. For the median comparisons, we corrected for bootstrap bias and used simulation-calibrated pointwise 95% confidence intervals and approximate two-sided tests, with adjustment for multiple comparisons. Full statistical procedures are given in the Supplementary Material, Methods S2.4. The decile distributions describe variation among species combinations, without tests of individual decile changes.

### Observed–unobserved comparisons

We then compared observed and unobserved pairs and triplets along the LUI gradient to investigate how coexistence mechanisms relate to species loss from local communities. For each plot-year census from 2008 to 2015, we evaluated all combinations composed entirely of observed species and all combinations containing observed partners and exactly one unobserved species. Observed species had positive cover in that census, while unobserved species had zero cover. We used the same 50-species pool in all three regions. For example, if A, B and C were observed species and D was unobserved, the observed-only pairs would be A–B, A–C and B–C, and the triplet would be A–B–C. The pairs containing D would be A–D, B–D and C–D, while the triplets would be A–B–D, A–C–D and B–C–D.

For each census, we calculated the percentage of feasible combinations and the mean niche and fitness differences within each group, separately for pairs and triplets, using the same fitted population models at the LUI recorded for that plot and year. We subtracted the unobserved-group summary from the observed-group summary, giving one difference per metric and combination size in each census, with feasibility differences expressed in percentage points. These analyses used 1,118 complete censuses from 150 plots within the analysed LUI range. We used Gaussian mixed-effects models to test how these differences varied with continuous LUI, accounting for region, year and repeated observations within plots. From these fitted models, we estimated the observed–unobserved difference at LUI 0.5 and 3 and the change between these endpoints. As fewer species were observed at high LUI, we also checked whether changing group sizes could bias these results by randomising species identities while retaining the number present in each census. Details of the statistical tests, randomisation check and graphical summaries are given in the Supplementary Material, Methods S3.

### Indirect effects across assemblage sizes

To evaluate whether indirect effects increased or reduced feasibility compared with pairs of species (where indirect interactions through other species cannot occur), we computed the community-pair differential (Fig. 1). Following Saavedra et al. (2017), we calculated the difference between the size of the full-assemblage feasibility domain and the domain in which all constituent pairs are simultaneously feasible in isolation, under the same growth- rate vector. Domain sizes were measured as probabilities over relative combinations of positive intrinsic growth rates, with fitted interaction coefficients held fixed. The differential ranges from −1 to 1, with positive values indicating greater opportunities for feasibility in the full assemblage and negative values indicating the opposite.

The differential compares the sizes of the two domains, but does not show how much they overlap. We therefore calculated community-pair overlap, which measures the proportion of the full-assemblage feasibility domain in which all constituent pairs are also feasible in isolation. Values close to zero indicate greater dependence on indirect interactions, whereas values close to one indicate less dependence. Overlap is undefined when the full feasibility domain is empty, so these assemblages were excluded from overlap means at each LUI.

Because the number of possible combinations increases rapidly with assemblage size, we sampled 1,000 combinations at each of four assemblage sizes (3, 5, 7 and 11 species) from the 50-species pool. We drew ten independent samples for each assemblage size to assess how the choice of species combinations affected the results. Within each sample, combinations were drawn uniformly without replacement and retained across all eleven LUI values from 0.5 to 3. We averaged both metrics across the assemblages in each sample.

We then assessed LUI responses using paired contrasts within each sample, comparing low (0.5), middle (1.75) and high (3) LUI. Middle minus low and high minus middle describe the two parts of the gradient, while high minus low measures net endpoint change. We also compared adjacent assemblage sizes at each of these LUI values. However, because assemblages were sampled separately at each size, these comparisons do not represent adding species to the same assemblage. We averaged contrasts across the ten samples, obtained 95% Monte Carlo intervals (Morris et al., 2019) and recorded how many samples agreed on the direction of each contrast. These intervals describe precision from sampling species combinations with the fitted interaction coefficients held fixed. Full details of the numerical procedures, graphical summaries and statistical comparisons are provided in the Supplementary Material, Methods S4.

### Analyses of individual LUI components

The LUI index combines management practices that can affect plant communities in different ways. To complement the overall LUI analysis, we repeated the population-model and structural analyses separately for fertilisation, mowing and grazing, replacing LUI with each component and retaining the model structure described above. We examined correlations among the practices across the full dataset and separately within each region (Table S1) to guide interpretation of the component associations (Dormann et al., 2013). All models used a common set of 1,135 annual transitions across 150 plots. We excluded *Brachypodium pinnatum* and *Thymus pulegioides* from these comparisons because each species occurred with positive cover and positive fertilisation in only one transition between two consecutive years. Structural analyses therefore used a common pool of 48 species, for which we evaluated all pairs and triplets.

We summarised structural niche and fitness differences with empirical medians and deciles across all combinations. For each component, we compared feasibility and the medians of these metrics between the minimum and maximum of its observed range. We used a joint parametric bootstrap to account for uncertainty in the fitted population parameters, as described in the Supplementary Material, Methods S1.1. We also repeated the indirect- effects analysis for assemblages of 3, 5, 7 and 11 species, using ten independent samples of 1,000 combinations per size from this common pool. Within each sample, the same combinations were evaluated across component values and across all three components. We assessed component-level and assemblage-size contrasts using the sampling-precision measures described above, with the minimum, midpoint and maximum of each component range as reference levels. Full details of the evaluation levels, uncertainty estimates and statistical comparisons are given in the Supplementary Material, Methods S5.

### Richness reconstruction and species-occurrence assessment

We then tested whether our models could reproduce the decline in richness observed along the LUI gradient. We compared reconstructed richness with maximum richness recorded within the same 50-species pool. For each plot-year census, we evaluated all pairs and triplets using the previously fitted population models at the LUI recorded for that plot and year. Each species was counted once if it participated in at least one feasible combination, separately for pairs and triplets. Triplets required feasibility of the full three-species assemblage. For instance, if A–B and C–D were feasible pairs, all four species would be counted, even if B–C was not feasible. Separate feasible combinations can occur in different patches, potentially allowing more species to coexist across a field than within an individual patch (García-Callejas et al., 2021). However, their union does not establish simultaneous feasibility of all counted species.

To examine whether facilitative effects allowed additional species to participate in feasible combinations, we compared two interaction scenarios. The first retained the fitted competition and facilitation. In the second, we replaced every positive interaction coefficient, including positive intraspecific effects, with a very small negative value (−10^−6^), while intrinsic growth rates and all other interaction coefficients remained unchanged. We also counted feasible pairs and triplets along the LUI gradient and recorded how many feasible triplets each species participated in under both scenarios.

We also asked whether species participating in more feasible combinations were those actually recorded in the field. We used fivefold cross-validation, dividing whole plots into five groups and assessing each group using population models fitted to the other four (Roberts et al., 2017). All records from the test plots were excluded from model fitting. At the LUI recorded for each test census, we ranked the 50 species by the fraction of their possible pairs or triplets that were feasible under each interaction scenario. Potential partners came from the full species pool, including locally absent species. As a benchmark, we also ranked species by how often they were recorded in the fitting plots from the same region. This provided a simple comparison based on which species were common in other fields.

We measured how well these rankings distinguished recorded from absent species using the area under the receiver operating characteristic curve (AUC) (Pearce & Ferrier, 2000). AUC measures how often a recorded species ranks above an absent species in the same census, without requiring a cut-off for predicting presence. A value of 0.5 indicates chance and 1 perfect discrimination. We summarised AUC overall and across LUI, compared each feasibility ranking with chance and the frequency benchmark, and compared the two interaction scenarios. Uncertainty intervals used whole-plot bootstrap resampling with fitted models held fixed. These analyses assessed species rankings rather than reconstructed richness or occurrence probabilities. Full procedures are described in the Supplementary Material, Methods S6.

Analyses were implemented in R (R Core Team, 2025). A full list of the software used can be found in the Supplementary Material, Software used. Computationally intensive analyses were run on the Hyperion cluster at the Donostia International Physics Center (DIPC).

## Results

Intraspecific interactions were predominantly competitive along the LUI gradient, whereas interspecific interactions included both competition and facilitation (Fig. S2A–B). Mean intraspecific effects became more negative between LUI values of 0.5 and 3 (change = −1.04 [95% CI: −1.70, −0.39], *z* = −3.08, *p* = 0.017), while mean interspecific effects showed no detectable change. The fitted mean intrinsic growth rate fell from 0.08 to −0.03, and the median from 0.06 to 0.02, between low and high LUI, although the intervals for both changes included zero (both *p* ≥ 0.489, Fig. S2C). Pairs and triplets containing at least one facilitative interspecific effect remained common throughout the gradient, with no detectable change in prevalence (both *p* = 0.356, Fig. S2D, Table S3).

Across all combinations in the 50 species pool, fitted feasibility declined from 36.8% to 29.1% for pairs and from 22.0% to 15.0% for triplets between LUI 0.5 and 3 (Fig. 2A). These declines were not statistically significant for either pairs (*z* = −0.86, *p* = 0.390) or triplets (*z* = −1.09, *p* = 0.277, Table S4). Median niche differences remained high across the gradient for both pairs and triplets. Pair median niche differences changed little, from 0.97 to 1.01, whereas triplet medians rose from 0.87 to 1.01 at LUI value of 2 before falling to 0.95 at the end of the LUI gradient (Fig. 3A). Fitted median fitness differences were larger at high LUI for both pairs and triplets (Fig. 3B). However, these changes in median niche and fitness differences were not statistically significant (Table S5). Both metrics also varied substantially among combinations throughout the gradient (Fig. S3).

**Figure 2.**
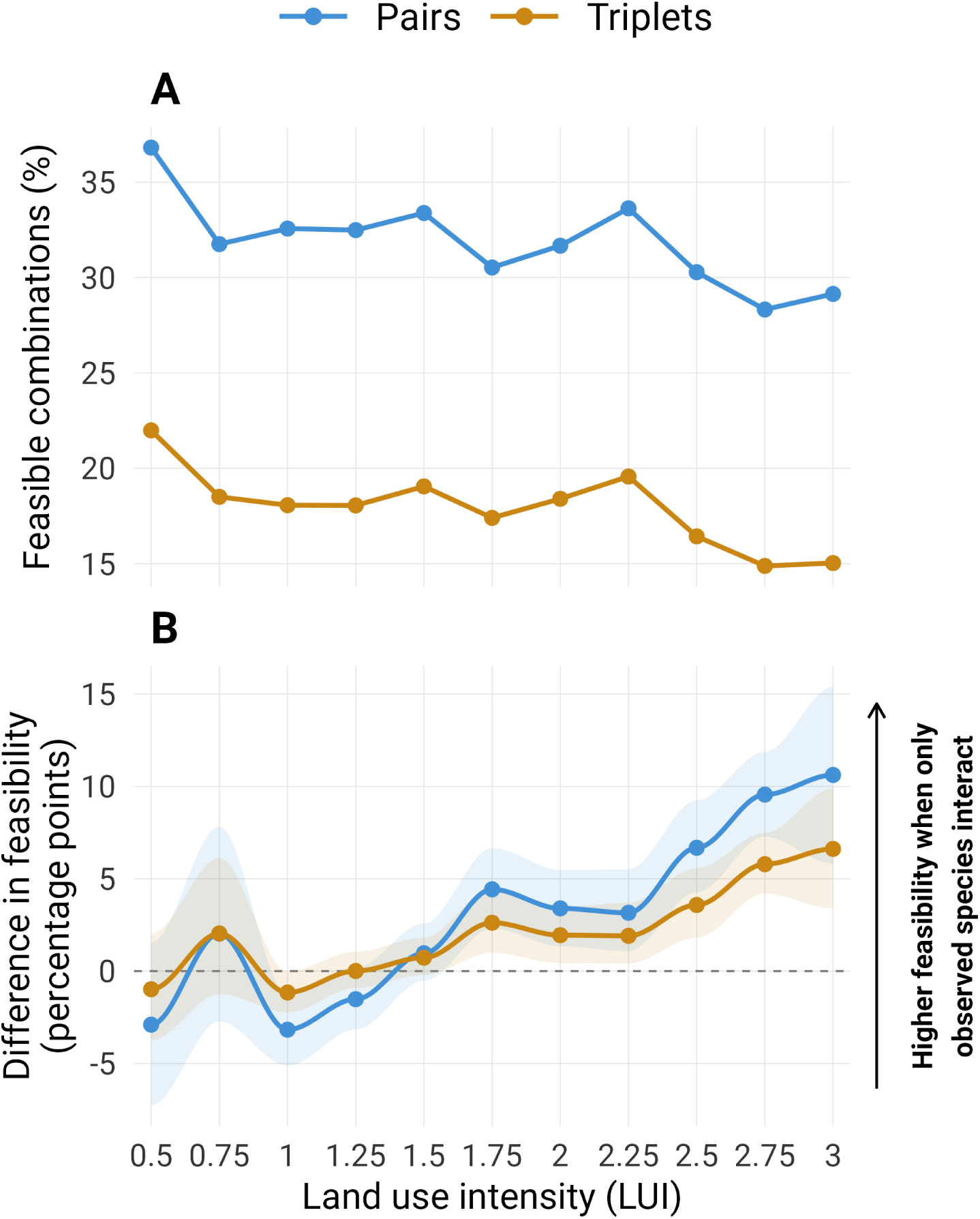
Feasibility across land use intensity (LUI) for pairs (blue) and triplets (orange). (A) Percentage of the 1,225 pairs and 19,600 triplets from the 50-species pool with positive equilibrium abundances for every member. (B) Difference between combinations composed entirely of locally observed species and combinations containing exactly one unobserved species and otherwise observed partners (observed minus unobserved, in percentage points). Census contrasts are averaged within plot and quarter-unit LUI bracket, then equally across plots. Shading shows pointwise 95% whole-plot bootstrap intervals, stratified by region; the dashed line marks zero. Points show calculated estimates, with interpolated curves and shading. Inference for A and B is reported in Table S4 and Table S6, respectively. Component-specific patterns are shown in Fig. S4.

**Figure 3.**
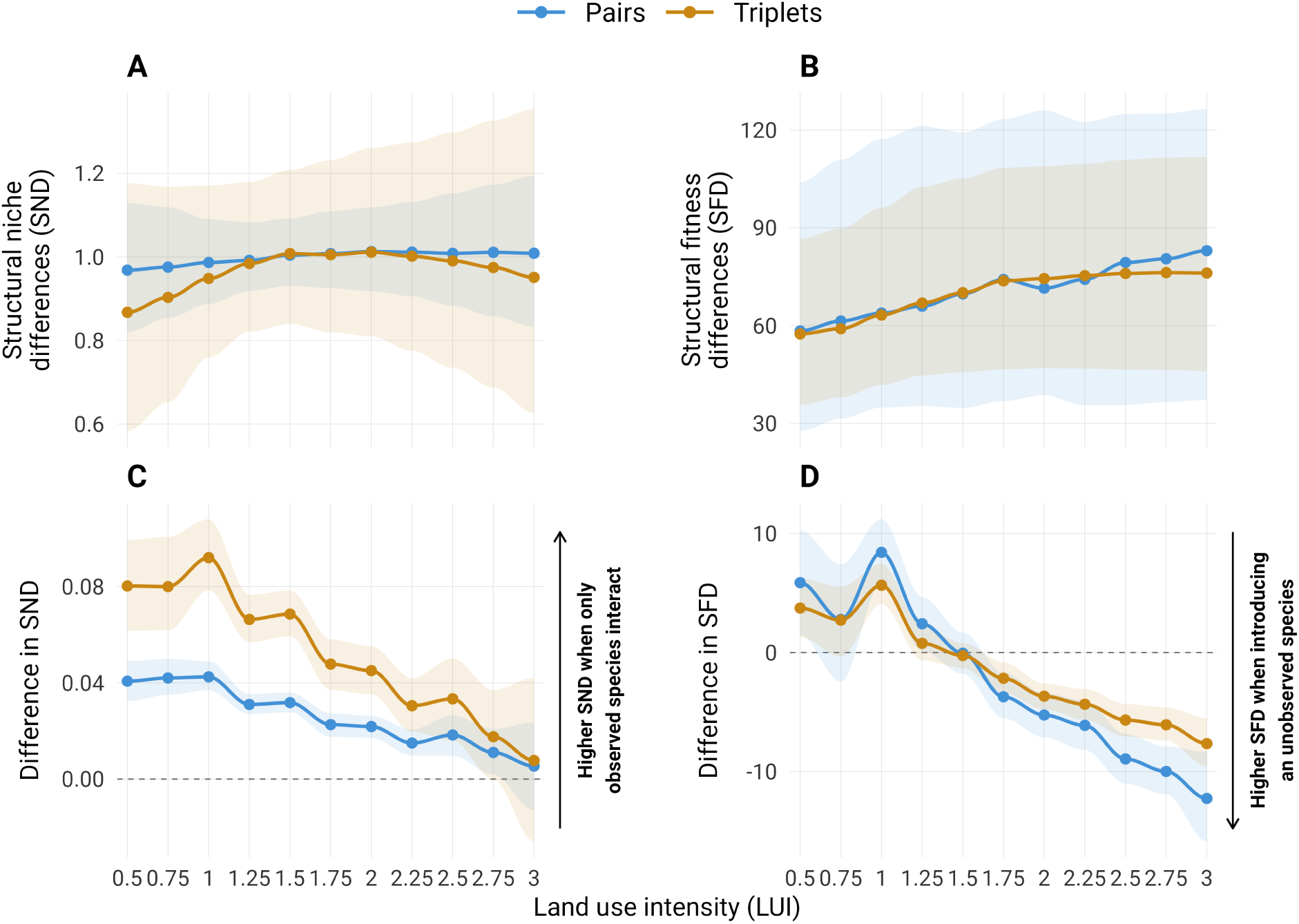
Structural niche differences (SND) and structural fitness differences (SFD) across land use intensity (LUI) for pairs (blue) and triplets (orange). (A, B) Medians across all combinations of the 50-species pool, with interquartile ranges. (C, D) Differences in census- level mean SND and SFD between entirely observed combinations and combinations containing exactly one unobserved species and otherwise observed partners (observed minus unobserved). Census contrasts are averaged within plot and LUI bracket, then equally across plots; shading shows pointwise 95% whole-plot bootstrap intervals. Dashed lines mark zero. Points show calculated estimates, with interpolated curves and shading. Pair and triplet values describe different geometries and are interpreted within assemblage size. Median contrasts across the species pool and observed–unobserved comparisons are reported in Table S5 and Table S7, respectively. Full quantile distributions and component- specific patterns are shown in Fig. S3 and Fig. S5.

To examine how these patterns related to local species occurrence, we compared combinations of species recorded together with alternatives containing one locally absent species. Observed combinations were less feasible than those containing an absent species at low LUI, but more feasible at high LUI (Fig. 2B). The observed–unobserved feasibility difference increased by 16.11 [95% CI: 13.60, 18.61] percentage points for pairs (*t*_960_ = 12.62, *p* < 0.001) and 8.18 [6.42, 9.94] for triplets (*t*_960_ = 9.14, *p* < 0.001, Table S6).

Observed combinations also had higher mean niche differences at both ends of the gradient, although this advantage became smaller as LUI increased (Fig. 3C). Between LUI 0.5 and 3, the niche-difference advantage decreased by 0.035 [95% CI: 0.028, 0.042] for pairs (*t*_960_ = −9.77, *p* < 0.001) and 0.072 [0.057, 0.087] for triplets (*t*_960_ = −9.53, *p* < 0.001, Table S7). Observed combinations had higher mean fitness differences at low LUI but lower fitness differences at high LUI than combinations containing an absent species (Fig. 3D). Between LUI 0.5 and 3, this shift amounted to a decrease in the observed–unobserved fitness difference of 23.99° [95% CI: 21.53, 26.44] for pairs (*t*_960_ = −19.18, *p* < 0.001) and 14.92° [13.41, 16.44] for triplets (*t*_960_ = −19.32, *p* < 0.001, Table S7). The randomisation check supported these changes along LUI, indicating that they were not simply due to the different numbers of observed and unobserved species (all six Holm-adjusted *p* ≤ 0.023, Supplementary Material, Methods S3.3).

The community-pair differential remained close to zero, although its pattern varied with assemblage size (Fig. 4A). It increased on average between LUI 0.5 and 3 for every assemblage size, while seven- and eleven-species assemblages had negative values at some intermediate LUI levels (Table S8 and Table S9). Community-pair overlap showed a more consistent pattern, with its highest values around LUI 1.5 and lower values in larger assemblages (Fig. 4B). Averaged across the ten samples, overlap at LUI 0.5, 1.75 and 3 was 0.83, 0.95 and 0.81 for triplets, compared with 0.03, 0.22 and 0.01 for eleven-species assemblages. All ten samples reproduced the increase from low to intermediate LUI, the subsequent decline towards high LUI and the lower overlap in each comparison between adjacent assemblage sizes (Table S8 and Table S9).

**Figure 4.**
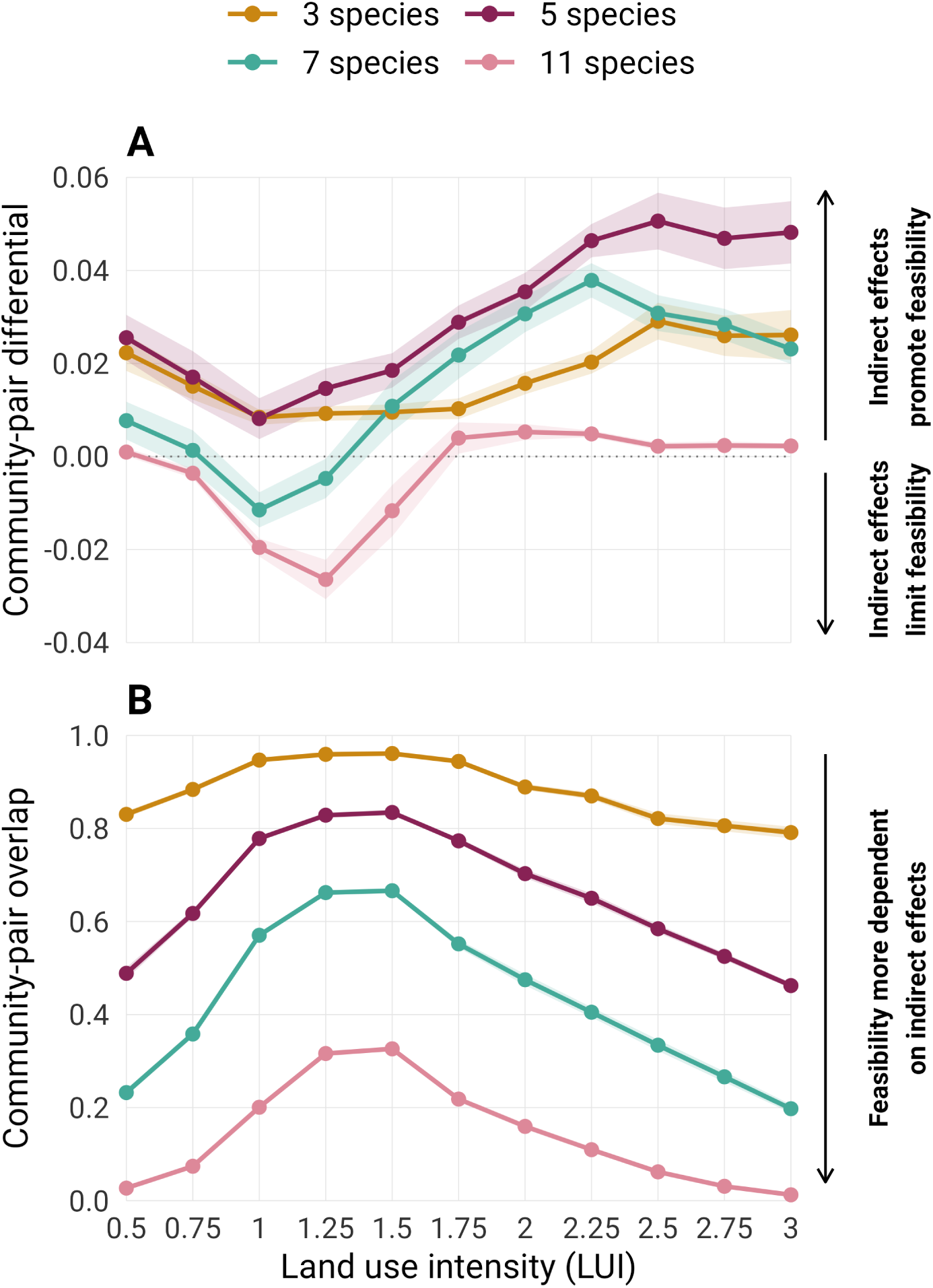
Indirect effects across land use intensity (LUI) for assemblages of 3, 5, 7 and 11 species from the 50-species pool. (A) Community-pair differential: the probability of full- assemblage feasibility minus that of simultaneous feasibility of all constituent pairs, over relative combinations of positive intrinsic growth rates. Positive and negative values indicate that indirect effects promote or limit feasibility, respectively. (B) Community-pair overlap: the proportion of the full-assemblage feasibility domain in which all constituent pairs are feasible. Lower overlap indicates greater dependence on indirect effects. Points show means for 1,000 sampled combinations per size, held constant across LUI; overlap means exclude empty full feasibility domains. Shading spans one standard deviation of the means from ten independently sampled sets of combinations. LUI contrasts and differences between assemblage sizes are reported in Table S8 and Table S9, respectively. See Fig. S6 for management-component comparisons.

When we examined fertilisation, mowing and grazing separately, feasibility showed no detectable change from the lowest to the highest level of any component (all *p* ≥ 0.610, Fig. S4, Table S10). Pairwise niche differences also changed little with any individual component, although fitted median values for triplets were lower at the upper end of each component gradient, decreasing from 0.99 to 0.86 under fertilisation, from 0.97 to 0.69 under mowing and from 0.97 to 0.72 under grazing. Fitted median fitness differences increased with fertilisation and decreased with mowing, but changed little under grazing. For both niche and fitness differences, the calibrated intervals for all median changes included zero (Fig. S5, Table S11). For indirect effects, overlap was lower at the upper end of each component gradient and in larger assemblages, with these directions reproduced in all ten samples, whereas the differential varied with the component and assemblage size (Fig. S6, Table S12 and Table S13).

Richness reconstructed from the fitted interaction matrices (with both competition and facilitation) exceeded the observed maximum throughout the LUI gradient (Fig. 5). Pair and triplet estimates declined initially following the observed richness decline but began to increase again at LUI 1.5 and 1.75, respectively. At the highest LUI, they averaged 39.7 and 50 species, compared with an observed maximum of 25. Eleven additional species participated in feasible triplets at LUI 1.75 compared with LUI 1.5, although the total number of feasible triplets fell (Fig. S7). Each of these species appeared in only 1–28 feasible triplets at LUI 1.75 (Fig. S8).

**Figure 5.**
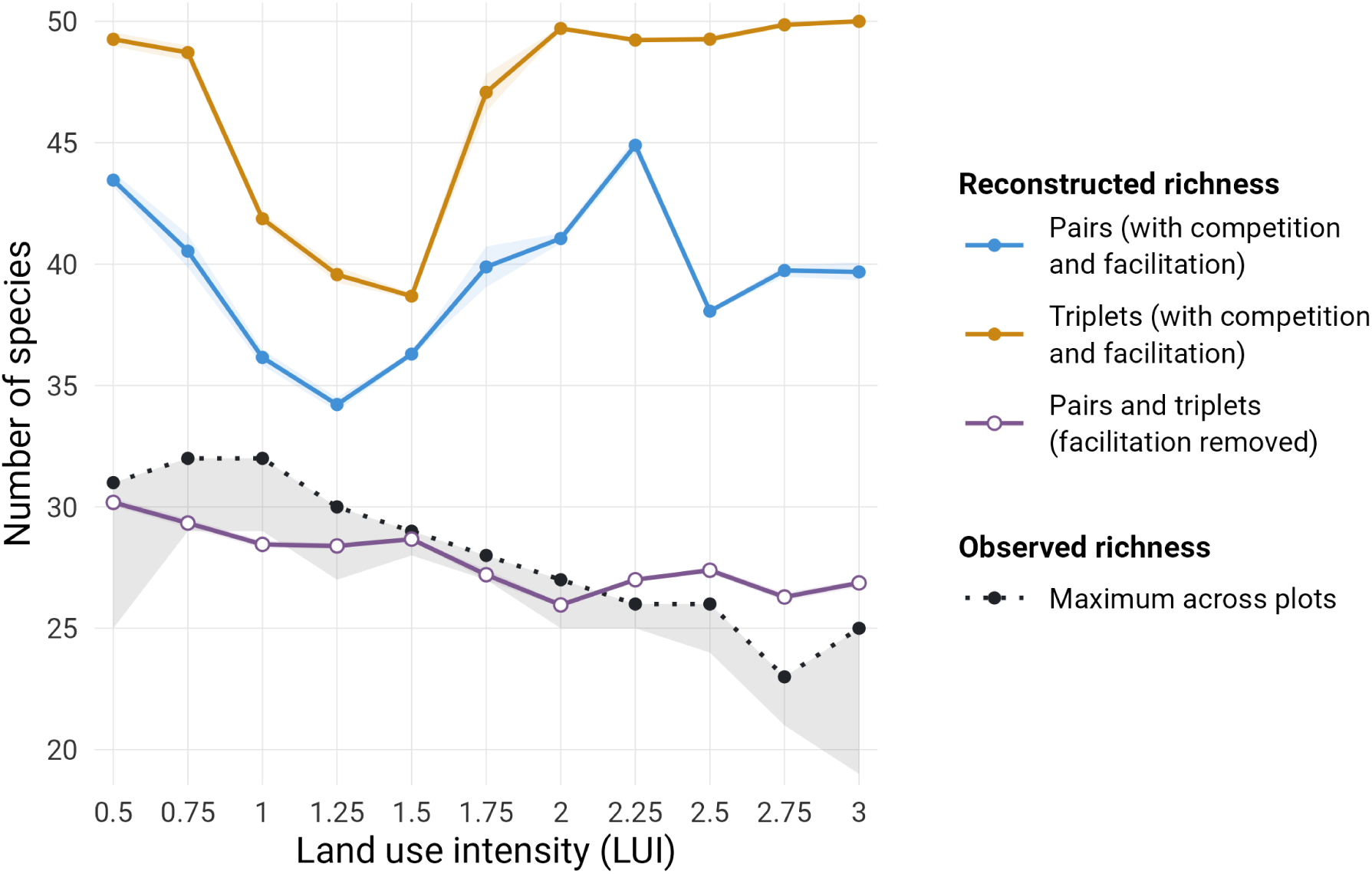
Observed maximum richness (dotted black) and richness reconstructed from feasible pairs (blue, filled circles) and triplets (orange, filled circles) with competition and facilitation, or with facilitation removed (purple, open circles), across land use intensity (LUI). Both scenarios evaluate the same 50-species pool at each census’s actual LUI and retain the fitted intrinsic growth rates. Facilitation removal replaces every positive fitted cover effect, including positive intraspecific effects, with −10^−6^; all non-positive effects remain unchanged. Each species is counted once if it belongs to at least one feasible combination, while triplets require feasibility of the full triplet. Reconstructions using pairs and triplets after facilitation removal produce the same richness results and are shown as one curve. Reconstruction means average censuses within plots and quarter-unit LUI brackets, then weight plots equally. Observed richness maxima are the largest counts across plots and censuses per LUI bracket. Shading shows pointwise 95% bootstrap intervals for reconstructed means and central 95% resampling ranges for observed maxima, using 9,999 whole-plot resamples within regions with fixed fitted coefficients. Participation in separate feasible combinations does not establish simultaneous feasibility of all counted species. Observed richness across the LUI gradient is shown in Fig. S1, while the number of feasible combinations at every LUI, species participation in those feasible combinations and discrimination between locally present and absent species are shown in Fig. S7, Fig. S8 and Fig. S9, respectively.

However, when facilitation was removed, reconstructed richness more closely followed the observed maximum and its decline with LUI. Pairs and triplets gave identical richness estimates, decreasing on average from 30.2 species in the lowest LUI bracket to 26.9 in the highest, compared with observed maxima of 31 and 25, respectively. The rebound at higher LUI observed when accounting for facilitation was removed when only considering competition. The main reason was because the eleven species responsible for the triplet rebound at LUI 1.75, all had negative fitted intrinsic growth rates and participated in no feasible pairs or triplets under this scenario.

We then assessed how well species’ participation in feasible combinations predicted their occurrence in the field. Overall, rankings based on feasibility were only slightly better than chance at distinguishing observed from absent species. Ranking species by how often they occurred in other plots in the same region performed much better, with an AUC of 0.837 [95% CI: 0.822, 0.851] (Fig. S9A). With competition and facilitation, mean AUC was 0.528 [0.516, 0.540] for pairs and 0.530 [0.518, 0.543] for triplets. These values changed little when facilitation was removed, to 0.533 [0.521, 0.546] for pairs and 0.535 [0.522, 0.547] for triplets, with no clear difference between the scenarios. In both scenarios, AUC was higher at high LUI, with point estimates around 0.47 in the lowest bracket and 0.59–0.61 in the highest (Fig. S9B–C).

## Discussion

Reversing losses in plant diversity following land use intensification requires a greater understanding of the mechanisms through which land use shapes local patterns of plant diversity. At high LUI, locally observed combinations were more feasible and had smaller fitness differences than alternatives containing locally absent species. This is consistent with intensification favouring local combinations of species with relatively similar fitness, even though fitness differences across the regional pool tended to increase. Niche differences remained high across the gradient, but the niche advantage of locally observed combinations weakened with intensification. Feasibility also depended more strongly on indirect interactions in larger assemblages, including under intensive management. Although facilitation expanded the set of species represented in feasible combinations, it biased the reconstruction of the decline of species richness with LUI. Removing facilitation brought reconstructed richness closer to the observed maximum and its decline with LUI. In this scenario, only species with positive fitted intrinsic growth rates belonged to feasible combinations. Together, these findings suggest that measures to mitigate diversity loss should consider how management practices affect the fitness of the species locally present, and the interactions among them.

Across the regional species pool, feasibility tended to decline with increasing LUI and fitted fitness differences increased. However, neither change was statistically significant. When we compared observed and unobserved combinations, we found that as LUI increased, observed combinations shifted from lower to higher feasibility and from larger to smaller fitness differences relative to combinations containing one locally absent species. This pattern is consistent with management favouring species with relatively similar fitness under the given environmental constraints. The larger fitness differences in combinations containing locally absent species suggest that fitness inequalities could contribute to species exclusion under intensive management. Our findings therefore agree with suggestions that species interactions can make some combinations from a regional pool more likely to occur than others (Medeiros et al., 2021). There are many processes affecting plant growth and interactions that could cause fitness differentiation. For instance, a soil-inoculation experiment found that microbes modified intrinsic growth rates and interspecific interactions, increasing fitness differences (Wang et al., 2024). In an experiment with perennial grassland species similar to those in our study, nutrient addition also increased fitness differences, particularly in combinations involving species disadvantaged by enrichment (Dostál, 2023). However, nutrient effects can differ among communities and treatments, as nitrogen enrichment reduced fitness differences in another grassland experiment (Granjel et al., 2023). It is plausible that fertilisation allowed nutrient-responsive plants to build taller canopies and capture a disproportionate share of light (DeMalach et al., 2017). Biomass removal through mowing or grazing may counter this shading and increase the light available to shorter neighbours, as shown by experiments manipulating herbivory and understorey light (Borer et al., 2014; Eskelinen et al., 2022). In our managed grasslands, maintaining biomass removal has been shown to support plant diversity during early reductions in management intensity, whereas the benefits of stopping fertilisation were less clear (Andraczek et al., 2023). Although the component estimates were uncertain and the management practices covaried, the opposing fitted fitness patterns under fertilisation and mowing are consistent with this idea.

In contrast to the pattern of fitness differences, median niche differences remained high across the gradient, with a peak at intermediate LUI in triplets. Niche differences were also higher in observed combinations than in unobserved alternatives, although this advantage diminished as LUI increased. Declining diversity was therefore accompanied by changes in both niche and fitness differences between observed combinations and potential alternatives. Intraspecific competition became stronger on average with LUI, while mean interspecific effects showed no corresponding shift. This stronger intraspecific competition is expected to limit growth as species become more abundant and could help maintain feasible combinations under intensive management (Adler, Smull, et al., 2018). A recent grassland experiment also linked larger structural niche differences to stronger intraspecific relative to interspecific interactions (Granjel et al., 2026). The processes underlying high niche differences might differ between low and high LUI grasslands. For instance, mycorrhizal richness increased with LUI in these grasslands (Gossner et al., 2016), and mycorrhizal fungi can promote plant coexistence through niche differentiation (Willing et al., 2024). At low LUI, trade-offs in nutrient use might play a greater role in maintaining niche differences, consistent with experiments showing that adding more limiting nutrients reduced grassland diversity (Harpole et al., 2016). Together, these findings suggest that the loss of plant diversity under intensive management need not reflect a uniform loss of niche differentiation.

Beyond pairwise effects, feasibility depended more strongly on indirect effects in larger assemblages, particularly at low and high LUI. These indirect interactions can weaken competition and make an assemblage feasible even when some pairs within it are not feasible on their own (Levine et al., 2017; Saavedra et al., 2017). For instance, native perennial plants have been shown to compete less strongly in multispecies groups than in pairs (Aschehoug & Callaway, 2015). In our analyses, indirect effects sometimes widened and sometimes narrowed the range of growth-rate combinations allowing feasibility. The average changes were small and varied with assemblage size and LUI. At the upper ends of the fertilisation, mowing and grazing gradients, feasibility also depended more strongly on indirect effects, although these effects did not consistently favour feasibility across assemblages of different sizes. Earlier work in our study system found that intensification reduced intransitivity and that greater intransitivity was associated with higher species richness (Soliveres et al., 2015). But intransitive loops are only one way that species can influence one another indirectly, so fewer loops do not necessarily mean weaker indirect effects. Moreover, greater intransitivity does not necessarily favour coexistence, as past work has shown that drought can reduce predicted coexistence among annual plants even when intransitivity remains similar or increases (Matías et al., 2018). Indirect effects therefore remained important under intensive management, but greater dependence on them did not necessarily mean a wider range of conditions allowing feasibility.

Identifying feasible species combinations did not fully reconstruct the richness observed in the field. Our reconstruction counted each species belonging to at least one feasible pair or triplet. With both competition and facilitation, reconstructed richness initially declined but then rebounded towards higher LUI, even as the total number of feasible combinations fell. This rebound was driven by species that appeared in only a few feasible combinations and all had negative fitted intrinsic growth rates. Removing facilitation excluded these species from feasible combinations and brought reconstructed richness closer to the observed maximum and its decline with LUI. In this scenario, reconstructed richness equalled the number of species with positive fitted intrinsic growth rates. These results also help explain why changes in niche and fitness differences did not translate directly into changes in richness.

Those metrics describe the conditions allowing particular combinations to be feasible (Saavedra et al., 2017), whereas reconstructed richness depends on which species are represented across feasible combinations. The feasibility of the species responsible for this rebound depended on positive neighbour effects, consistent with theory showing how facilitation can offset negative intrinsic growth and allow otherwise infeasible combinations (Godoy et al., 2026; Hart, 2023). Positive interspecific effects were common throughout the LUI gradient. For example, *Holcus lanatus* continued to occur in these grasslands despite fitted intrinsic growth rates ranging from −0.17 to −0.35 between LUI 1.75 and 3. Positive fitted effects from *Lolium perenne* and *Plantago lanceolata* throughout this range could offset the negative fitted intrinsic growth of *H. lanatus* in the model. This possible role for facilitation is consistent with evidence that it can operate beyond conspicuously harsh environments (Liancourt & Dolezal, 2021). Whether these benefits support coexistence depends on how they combine with competition and the processes that limit population growth (Hart, 2023; Hart & Marshall, 2013). The closer agreement with observed maximum richness after removing facilitation raises the possibility that positive effects were overestimated or that the conditions required for their benefits are not always met in the field. Experiments measuring growth with and without neighbours could test whether *H. lanatus* and other species with negative fitted intrinsic growth rates depend on facilitation under intensive management. Our results therefore reinforce the value of considering how facilitation and intraspecific competition together shape feasibility in human-modified ecosystems (James et al., 2025; Losapio et al., 2021).

Although removing facilitation improved agreement with observed maximum richness, it was not accompanied by a clear improvement in distinguishing locally present from absent species. Rankings based on species feasibility performed only slightly better than chance overall and worse than rankings based on how often species occurred in other plots, although performance increased at high LUI. Several biological processes could contribute to this mismatch. Combinations can be infeasible because of dominant species that are predicted to exclude their neighbours but are actually observed in the field (García-Callejas et al., 2021). Conversely, species may lack suitable partners or conditions locally, despite having feasible combinations within the regional pool (HilleRisLambers et al., 2012). Separate feasible combinations may also occur in different patches, so pooling their species does not identify a single local community (García-Callejas et al., 2021). Equilibrium predictions may also miss opportunities for coexistence created by changing conditions. For example, rainfall fluctuations promoted coexistence in an annual grassland because periodic early-season drought reduced competition from a dominant grass and allowed a forb to recover from low abundance (Hallett et al., 2019). These possibilities illustrate why a positive equilibrium does not establish that a combination will occur in the field (Clark et al., 2026), and why field predictions require separate validation (Tredennick et al., 2021).

Several of our findings also suggest non-linear changes in opportunities for coexistence around intermediate LUI. Dependence on indirect interactions was lowest around LUI 1.5, while median niche differences in triplets peaked near LUI 2. Locally observed combinations also gained advantages in feasibility and fitness similarity over unobserved alternatives in this part of the gradient, where richness reconstructed with the fitted interactions began to rebound. Abrupt transitions in diversity and ecosystem functioning have also been documented in these grasslands, with thresholds varying among responses (Saiz et al., 2026). Functional trait distributions also shifted abruptly with increasing LUI, indicating changes in the balance of plant resource-use strategies (Le Bagousse-Pinguet et al., 2025). These findings raise the possibility that changes in species interactions accompany broader shifts in grassland diversity and functioning. Understanding in future work how these changes are connected could help explain why the consequences of further intensification differ along the management gradient.

Our work also has limitations that future studies should address to improve predictions. Growth rates and interactions were inferred from changes in plant cover without separately modelling dispersal or measurement error. Additional measurements could help distinguish their effects from population growth. In addition, intrinsic growth rates describe hypothetical growth at zero cover of the modelled species and have not been independently measured. Negative fitted rates therefore do not imply that a species is rare in the field, as some were common under intensive management. These estimates can guide experiments measuring growth alone and with neighbours to test the inferred dependence on facilitation (Adler, Kleinhesselink, et al., 2018). Moreover, data support varied among species and combinations of neighbours, with fewer observations for some at high LUI. Targeted measurements under intensive management would help evaluate predictions that extend beyond a species’ observed LUI range (Conn et al., 2015). Our fitted population model assumes linear, additive effects of cover on per capita log growth. Comparing alternative forms of density dependence and higher-order models would help assess how strongly the conclusions depend on this formulation (Letten & Stouffer, 2019; Terry & Armitage, 2024). For indirect effects, repeated samples assessed how results changed with the species included but did not capture uncertainty in the fitted growth rates and interactions. Our analyses were also restricted to the 50 most common species, which showed a weaker decline in observed richness than the full set of recorded species (Fig. S1). Rarer species can be particularly sensitive to management in these grasslands (Allan et al., 2014), but their sparse records limit reliable estimation of growth rates and interactions. Finally, positive equilibrium abundances do not guarantee persistence through time (Clark et al., 2026). Assembly graphs could help connect these feasibility results with possible sequences of species establishment and loss (Godoy et al., 2024; Song, 2025), once population dynamics and the success of arriving species have been assessed.

Our study suggests that land use intensification favours fitness similarity among species found together locally as plant diversity declines. Niche differences remained high across the gradient, although the niche advantage of locally observed combinations weakened with intensification. Indirect interactions continued to shape feasibility under intensive management. The facilitation comparison also suggests that some species may depend on their neighbours. These results have implications for restoring managed grasslands. If the remaining species at high LUI form feasible combinations more often with one another than with potential newcomers from the regional pool, restoring propagule supply alone may not be sufficient to recover richness while intensive management continues. Experiments introducing locally absent species under contrasting management regimes and with different resident neighbours could test whether fitness differences restrict establishment and whether reducing management intensity allows these species to establish. More broadly, using time series to estimate growth rates, species interactions and coexistence mechanisms could help develop a more mechanistic understanding of biodiversity responses to global change and guide experiments to test the processes involved.

## Author contributions

OG, FvP, SS and EA conceptualised the study. RRG, OG and EA conducted the statistical analyses. RRG prepared the figures. RRG, OG and EA wrote the manuscript with substantial contributions from all authors.

## Acknowledgements

We thank Serguei Saavedra for his guidance on the correspondence between the population and statistical models. We acknowledge the computational resources and technical support provided by the Donostia International Physics Center (DIPC), including access to the Hyperion cluster.

## Data availability statement

The data and R code supporting this study, including processed outputs for computationally intensive analyses, are available at https://anonymous.4open.science/r/lui-feasibility-jecol/. The original vegetation and land-use data are available through BExIS and Vogt et al. (2019), respectively.

