## Supplementary Material for "Land use intensification favours fitness similarity as plant diversity declines"

[1]Basque Centre for Climate Change (BC3), 48940 Leioa, Spain; [2]Doñana Biological Station (EBD-CSIC), Americo Vespucio 26, 41092 Sevilla, Spain; [3]Plant Ecology and Nature Conservation Group, Wageningen University, P.O. Box 47, 6700 AA Wageningen, The Netherlands; [4]Systematic Botany and Functional Biodiversity, Institute of Biology, Leipzig University, Johannisallee 21–23, 04103 Leipzig, Germany; [5]Department of Ecology, University of Alicante, Spain; [6]Environmental Studies Institute "Ramón Margalef", University of Alicante, Spain; [7]Institute of Plant Sciences, University of Bern, Altenbergrain 21, 3013 Bern, Switzerland; [8]Departamento de Ciencias Agrarias y del Medio Natural, Instituto Universitario de Investigación en Ciencias Ambientales de Aragón (IUCA), Universidad de Zaragoza, Spain; [9]Institute of Landscape Ecology, University of Münster, Heisenbergstr. 2, 48149 Münster, Germany. [†]Deceased March 2026.

### Contents


Supplementary Methods ... 3
- S1. Population-model uncertainty and parameter comparisons ... 3
- S2. Feasibility, structural metrics and regional comparisons ... 3
- S3. Observed–unobserved comparisons ... 5
- S4. Indirect effects across assemblage sizes ... 7
- S5. Individual LUI components ... 8
- S6. Richness reconstruction and species-occurrence assessment ... 9

Supplementary figures and tables ... 12

- Figures ... 12
  - Figure S1 LUI and observed richness ... 12
  - Figure S2 Population parameters and facilitation ... 13
  - Figure S3 SND and SFD quantiles ... 14
  - Figure S4 Feasibility across LUI components ... 15
  - Figure S5 SND and SFD across LUI components ... 16
  - Figure S6 Indirect effects across LUI components ... 17
  - Figure S7 Numbers of feasible combinations ... 18
  - Figure S8 Species participation in feasible triplets ... 19
  - Figure S9 Species-identity discrimination ... 20
- Tables ... 21
  - Table S1 Correlations among LUI components ... 21
  - Table S2 Species pool and occurrence frequency ... 22
  - Table S3 Population-parameter and facilitation contrasts ... 23
  - Table S4 Feasibility contrasts across the species pool ... 24
  - Table S5 Median SND and SFD contrasts across the species pool ... 25
  - Table S6 Observed-minus-unobserved feasibility ... 26
  - Table S7 Observed-minus-unobserved SND and SFD ... 27
  - Table S8 Indirect effects: LUI contrasts ... 28
  - Table S9 Indirect effects: assemblage-size contrasts ... 29
  - Table S10 Feasibility contrasts across LUI components ... 30
  - Table S11 Median SND and SFD: LUI components ... 31
  - Table S12 Indirect effects: component contrasts ... 32
  - Table S13 Indirect effects: size contrasts by component ... 35

Software used ... 37

Supplementary references ... 38

### Supplementary Methods

#### S1. Population-model uncertainty and parameter comparisons

##### S1.1. Population-model uncertainty

We used a joint Gaussian parametric bootstrap to carry uncertainty in the fitted population parameters into the structural analyses. Observed cover predictors, management values and missing-data patterns were held fixed. Simulated responses retained the fitted regional and plot random-intercept variances and temporal AR(1) covariance. They also included a working cross-species correlation estimated from marginal residuals after removing their species-specific covariance structure. Correlated Gaussian innovations were returned to each species' covariance scale and added to the fitted fixed-effect predictions. We then refitted the population models and recalculated the required summaries. Each bootstrap retained the cover predictors used to fit the corresponding population models. Feasibility and structural metrics were calculated for the selected 50-species pool, or 48 species in the component analyses. The main LUI analysis produced 998 successful replicates from 999 attempts.

##### S1.2. Population-parameter comparisons

We assessed changes between LUI 0.5 and 3 in the means and medians of interspecific effects, intraspecific effects and intrinsic growth rates. We also assessed the change in facilitation prevalence for pairs and triplets. A combination contributed to facilitation prevalence if it contained at least one positive directed interspecific cover effect, with intraspecific effects excluded from this measure.

These eight endpoint contrasts used pointwise 95% bootstrap percentile intervals and approximate two-sided Wald tests based on bootstrap standard errors. We applied the Benjamini–Hochberg adjustment across the eight tests. Two further Wald tests assessed changes in mean interspecific and intraspecific effects jointly, and changes in pair and triplet facilitation prevalence jointly. These used the bootstrap covariance, two degrees of freedom, and a separate Benjamini–Hochberg adjustment across the two tests. Endpoint contrasts describe net changes across the analysed range, rather than testing a monotonic response throughout it.

#### S2. Feasibility, structural metrics and regional comparisons

For an assemblage of *n* species at a given LUI, $\mathbf{r}$ contains its fitted intrinsic growth rates and $A = \left(\alpha_{ij}\right)$ its interaction coefficients. The interaction coefficients follow the population model

(Equation 2). Positive $\alpha_{ij}$ denotes facilitation and negative $\alpha_{ij}$ competition. Structural niche and fitness differences follow Saavedra et al. (2017).

##### S2.1. Feasibility

For an invertible interaction matrix, equilibrium relative covers and the feasibility domain are

$$\mathbf{N}^{*} = -A^{-1}\mathbf{r}, \qquad \mathcal{F}(A) = \left\{\mathbf{r} : -A^{-1}\mathbf{r} > 0\right\}. \tag{S1}$$

A combination is feasible only if every equilibrium cover is strictly positive. This condition concerns the existence of a positive equilibrium, but does not establish its dynamical stability (Grilli et al., 2017). Combinations with undefined equilibria are not counted as verified feasible.

##### S2.2. Structural niche differences

Structural niche differences (SND) measure the normalised angular size of the feasibility domain:

$$\Omega(A) = 2^{n}\,\Pr\left(-A^{-1}\mathbf{u} > 0\right), \tag{S2}$$

where **u** has a uniformly distributed direction over the whole unit sphere. Larger values indicate a wider range of feasible growth-rate directions. The normalisation makes SND equal to one for self-limited species without interspecific interactions. With facilitation, values can exceed one. Niche differences are dimensionless and are interpreted within assemblage size. For pairs and triplets, we calculated the angular probability analytically using the equivalent centred-Gaussian orthant probability.

##### S2.3. Structural fitness differences

Structural fitness differences (SFD) measure the angle between the fitted growth-rate vector and the centroid of the feasibility domain, $\mathbf{r}_c$:

$$\mathbf{r}_c = -\frac{1}{n}\sum_{j=1}^{n}\frac{\boldsymbol{\alpha}_j}{\left\|\boldsymbol{\alpha}_j\right\|}, \qquad \theta = \frac{180}{\pi}\arccos\left(\frac{\mathbf{r}\cdot\mathbf{r}_c}{\|\mathbf{r}\|\,\|\mathbf{r}_c\|}\right). \tag{S3}$$

Here, $\boldsymbol{\alpha}_j$ is column $j$ of $A$ and $\|\cdot\|$ denotes vector length. To find the centroid, we give the vectors defining the domain's boundaries a length of one and then average them. Fitness differences are expressed in degrees from 0 to 180. Zero indicates alignment with this centroid, which does not necessarily mean equal growth rates or equal fitness. We assess feasibility directly from equilibrium covers, rather than comparing SND numerically with SFD.

##### S2.4. Regional contrasts and calibration

Feasibility contrasts compared the percentage of feasible combinations at LUI 3 with that at 0.5. They used pointwise 95% percentile intervals and approximate two-sided Wald tests based on the population-model bootstrap described in S1.1, without a multiple-testing adjustment.

We summarised niche and fitness differences with empirical medians, interquartile ranges and deciles across combinations. No quantile regressions were fitted. We compared medians between LUI 0.5, 1.75 and 3. For each contrast, estimated bias was the mean bootstrap contrast minus the fitted contrast. We subtracted this bias from the fitted contrast and used the bootstrap standard deviation as its standard error.

To calibrate intervals and tests, we simulated additional datasets from the fitted population models. Each dataset was refitted, with 199 inner bootstrap refits attempted to estimate its contrast bias and standard error. Calibration scores measured the absolute error of the bias-corrected contrast relative to its generating-model value, divided by its estimated standard error. Of these datasets, 750 supplied calibration scores and a separate 250 assessed interval coverage. Pointwise 95% intervals used the 714th ordered calibration score as their critical value. Approximate two-sided test probabilities were $(b + 1)/751$, where $b$ was the number of calibration scores at least as large as the observed standardised contrast. We applied Holm adjustment across the 12 comparisons: two metrics, two assemblage sizes and three contrasts. Calibration describes uncertainty under the fitted models, whereas decile bands describe differences among species combinations.

#### S3. Observed–unobserved comparisons

##### S3.1. Census contrasts and graphical summaries

Using the same fitted population models, we evaluated pairs and triplets from the 50-species pool at the LUI recorded in each census. We compared combinations formed entirely by observed species with those containing observed partners and exactly one unobserved species. For each group, we calculated the percentage of feasible combinations and the mean niche and fitness differences, separately for pairs and triplets. We reported the differences as observed minus unobserved, with feasibility differences expressed in percentage points.

For the figures, we grouped censuses into LUI brackets of width 0.25, centred at 0.5, 0.75, …, 3. Each bracket included its lower boundary and excluded its upper boundary, giving an overall range from 0.375 to below 3.125. The analysis included 1,118 censuses from 150 plots with complete species-presence data within this range. Within each bracket, we averaged census differences within each plot and then averaged the plot means, giving every contributing plot equal weight. Pointwise 95% percentile intervals used 999 whole-plot bootstrap resamples within regions, retaining all censuses from each selected plot (Field & Welsh, 2007). The fitted population-model estimates were held fixed during resampling.

**S3.2. Statistical comparisons**

We fitted six Gaussian mixed-effects models to the census-level differences, analysing each metric separately for pairs and triplets. For these tests, we used each census's group difference and recorded LUI, rather than the bracket summaries shown in the figures. Continuous LUI was centred at 1.75. Models included categorical region and year effects, a plot random intercept, temporal AR(1) residual correlation within plots and region-specific residual variances. They were fitted by restricted maximum likelihood (REML).

We estimated the observed-minus-unobserved difference at LUI 0.5 and 3 and the change between these endpoints, averaging predictions equally across regions and years. Two-sided $t$ tests and pointwise 95% $t$ intervals used containment degrees of freedom. Holm adjustment covered all 18 comparisons. As a check on the assumed response shape, we repeated the endpoint comparisons after including a quadratic LUI term, with a separate adjustment across that set of 18 tests. The fitted population-model estimates were held fixed throughout the statistical comparisons and graphical resampling.

**S3.3. Checking the effect of changing species numbers**

We used 9,999 randomisations to assess whether the changing numbers of observed and unobserved species could explain the observed–unobserved contrasts. In each randomisation, we permuted the labels of the 50 species independently within each plot, using the same mapping across all its censuses. This retained each census's richness and LUI, the numbers of possible combinations in both groups, and the overlap in species membership between years. Fitted population parameters were held fixed, and the three contrasts were recalculated at each census's recorded LUI. This check addressed species numbers rather than differences in species frequencies or regional availability.

To summarise each empirical and randomised dataset in the same way, we used a linear model with LUI and categorical region and year effects, weighting censuses so that each plot contributed equally. We estimated contrasts at LUI 0.5 and 3 and their difference, averaging regions and years equally. These provided separate randomisation statistics rather than refitting the mixed-effects models in S3.2. Two-sided probabilities were calculated as $(1 + b)/10,000$, where $b$ was the number of randomised contrasts at least as large in absolute value as the empirical contrast. Holm adjustment covered all 18 comparisons, comprising the two endpoints and their change for three metrics and two assemblage sizes.

#### S4. Indirect effects across assemblage sizes

##### S4.1. Community-pair differential and overlap

Let $F$ denote feasibility of the full assemblage and $P$ simultaneous feasibility of every constituent pair considered in isolation. Both events are evaluated under the same growth-rate vector. For directions drawn uniformly from the positive unit sphere, define

$$p_F = \Pr(F), \qquad p_P = \Pr(P), \qquad D = p_F - p_P, \qquad O = \Pr(P \mid F) = \frac{\Pr(F \cap P)}{p_F}. \tag{S4}$$

The community-pair differential, $D$, ranges from −1 to 1. Positive values mean that more growth-rate directions permit full-assemblage feasibility than simultaneous pairwise feasibility, whereas negative values indicate the reverse. Community-pair overlap, $O$, is the proportion of the full-assemblage feasibility domain in which all constituent pairs are also feasible in isolation. Lower overlap indicates greater dependence on indirect interactions. An overlap of one means that the full-assemblage domain is contained within the pairwise domain, rather than establishing that the two domains are identical.

Both metrics concern positive intrinsic growth rates, unlike the whole-sphere definition of niche differences, which can include negative rates when facilitation permits feasibility. Overlap is undefined when the full-assemblage domain is empty. Those assemblages were excluded from overlap means separately at each LUI, while a nonempty full domain with an empty intersection has zero overlap.

##### S4.2. Assemblage sampling and differential

We drew ten independent samples of 1,000 assemblages at each size of 3, 5, 7 and 11 species from the 50-species pool. Assemblages were sampled uniformly without replacement within each sample, and the same combinations were retained across all eleven LUI values from 0.5 to 3 in steps of 0.25. The fitted interaction coefficients at each LUI were held fixed throughout these calculations. For each assemblage and LUI, we estimated the differential using 8,192 independent directions drawn uniformly from the positive unit sphere by normalising absolute Gaussian draws. Full-assemblage and simultaneous pairwise feasibility were assessed using the same direction.

##### S4.3. Estimating community-pair overlap

We estimated overlap by sampling directly within the full-assemblage feasibility domain. This avoids relying on the ratio of two small numbers of feasible draws when that domain is narrow. Linear programming first checked whether the full domain and its intersection with the pairwise domain were empty. Nonempty domains were sampled using a constrained-Gaussian method with two chains and adaptive sampling. Checks required split $\hat{R} \leq 1.05$, effective sample size of at least 200 and Monte Carlo standard error no greater than 0.025. Independent rejection sampling was used where required.

##### S4.4. Contrasts and sampling precision

We averaged each metric across assemblages within each sample. Curves show the first sample's means, with ribbons extending one standard deviation of the ten sample means above and below them. Within each sample, we calculated middle minus low, high minus middle and high minus low contrasts, using LUI 0.5, 1.75 and 3 as the low, middle and high levels, respectively. We also compared adjacent assemblage sizes at each level as 5 minus 3, 7 minus 5 and 11 minus 7 species. Because different assemblages were sampled at each size, size contrasts do not represent adding species to the same assemblage.

Contrasts were averaged across the ten samples. Monte Carlo standard errors were their standard deviation divided by $\sqrt{10}$, and pointwise 95% Monte Carlo intervals used a $t$ distribution with nine degrees of freedom (Morris et al., 2019). We also recorded how many samples showed the same direction of change as the mean contrast. These measures describe uncertainty from sampling species combinations with interaction coefficients held fixed and do not include uncertainty in the fitted population parameters.

#### S5. Individual LUI components

##### S5.1. Models, species pool and evaluation levels

We replaced LUI with fertilisation, mowing or grazing and otherwise retained the population-model structure. All component fits used a common set of 1,135 annual transitions across 150 plots. One transition used in the composite-LUI analysis lacked a mowing record and was excluded from this common dataset. The structural analyses included 48 taxa. *Brachypodium pinnatum* and *Thymus pulegioides* were excluded from the component analyses because each had positive cover with positive fertilisation in only one transition.

The common 48-species structural pool contained 1,128 pairs and 17,296 triplets. We evaluated components in their original units over their observed ranges: 0–345.35 kg N ha$^{-1}$ year$^{-1}$ for fertilisation, 0–4 cuts year$^{-1}$ for mowing, and 0–1,644.167 livestock-unit days ha$^{-1}$ year$^{-1}$ for grazing. Fertilisation and grazing were evaluated at eleven equally spaced levels, and mowing at the five integer levels.

##### S5.2. Uncertainty and comparisons

Each component produced 999 successful joint population-model bootstrap replicates, with observed predictors fixed and models refitted as described in S1.1. Endpoint feasibility contrasts compared the maximum with the minimum of each component range. They used pointwise 95% percentile intervals and approximate two-sided Wald tests based on the bootstrap standard error of the change, without adjustment for multiple comparisons. Contrasts in median niche and fitness differences used the same endpoints, with bootstrap bias correction and simulation calibration as described in S2.4. For these median comparisons, there were four contrasts per component (two metrics and two assemblage sizes), giving 12 comparisons across the three components.

Each component had 750 calibration datasets and 250 additional datasets for coverage assessment, with 199 inner bootstrap refits attempted per dataset. A coverage diagnostic led to a more conservative rule accounting for finite simulation precision across the 12 component contrasts. Pointwise intervals used the 728th ordered calibration score. Tests used one-sided upper binomial bounds on the calibration tail probabilities, calculated at confidence level $1 - 0.05/12$, followed by Holm adjustment across all 12 component comparisons. This safeguard concerns precision of calibration, rather than simultaneous coverage of all ecological effects. Coverage measured on the retained diagnostic datasets after refining the rule is not a fresh independent validation.

##### S5.3. Indirect effects and management correlations

For indirect effects, we used the procedures in S4, drawing ten independent samples of 1,000 assemblages at each size of 3, 5, 7 and 11 species from the common 48-species pool. Within each sample, combinations were retained across all levels and all three components. Contrasts used the minimum, midpoint and maximum of each component's range.

We described correlations among the three practices using Pearson's $r$ and Spearman's $\rho$, pooled across regions and separately within each region. Pointwise 95% percentile intervals used 1,999 whole-plot bootstrap resamples within regions, retaining all annual transitions from each selected plot. Approximate two-sided Wald tests used Fisher-transformed correlations and their bootstrap standard errors, with Benjamini–Hochberg adjustment across the 24 tests. Separate component fits describe management associations and do not identify independent effects of correlated practices (Dormann et al., 2013).

#### S6. Richness reconstruction and species-occurrence assessment

##### S6.1. Interaction scenarios

These interaction scenarios were used for richness reconstruction and species-occurrence assessment. The preceding analyses retained the fitted competition and facilitation. Both scenarios used the same 50-species pool, evaluating all 1,225 pairs and 19,600 triplets at each census's actual LUI. The first retained the fitted competition and facilitation. In the second, every strictly positive interaction coefficient ($\alpha_{ij} > 0$) was replaced with $-10^{-6}$ after evaluating its LUI response. This included positive intraspecific effects. Intrinsic growth rates and non-positive interaction coefficients were unchanged. This replacement was applied to the fitted coefficients, without refitting models or filtering species.

##### S6.2. Richness and species participation

A species was counted once if it belonged to at least one pair or full triplet with strictly positive equilibrium covers for all members. Pairs and triplets were evaluated separately. Their union describes the species represented in feasible combinations but does not establish simultaneous feasibility of all counted species (García-Callejas et al., 2021). Exact duplicated rows or columns and other undefined interaction matrices were recorded and contributed no verified feasible participation.

The reconstruction used the same 1,118 censuses from 150 plots and quarter-unit LUI brackets as the local comparison. Reconstructed richness was averaged within plots and LUI brackets, then equally across contributing plots. Observed maximum richness was the largest single-census count in each bracket, restricted to the same 50-species pool. Pointwise 95% percentile intervals for mean reconstructed richness used 9,999 whole-plot bootstrap resamples within regions, retaining repeated censuses and holding the fitted coefficients fixed. For observed maxima, we reported the central 95% range of resampled maxima. This describes sensitivity to the sampled plots, rather than uncertainty about an unknown population maximum.

To describe how species enter these reconstructions, we also counted feasible pairs and triplets at each census's actual LUI (Fig. S7) and each species' participation in feasible triplets at the eleven exact LUI values (Fig. S8). The former used the same bracket summaries and bootstrap resamples as the richness curves. Participation counts describe combinations, rather than equilibrium abundance.

##### S6.3. Species-occurrence assessment

We used fivefold cross-validation stratified by region, with 120 training plots and 30 test plots per fold. All censuses from each test plot were excluded during population-model fitting (Roberts et al., 2017). The assessment was conditional on the 50-species target pool, which had been selected before cross-validation. Both scenarios used the corresponding training fits, applying the same post-fitting replacement of positive effects for facilitation removal.

At each test census's LUI, we ranked the 50 species by the fraction of their 49 possible pairs or 1,176 possible triplets that were verified feasible. Potential partners came from the whole target pool, regardless of their presence in the test census. Undefined combinations remained in these denominators and contributed no verified participation. As a benchmark, we ranked species by occurrence frequency in training plots from the test plot's region, averaging censuses within training plots and then weighting plots equally. This benchmark was regional, rather than restricted to the test census's LUI bracket.

AUC (area under the receiver operating characteristic curve) was calculated within each test census as the probability that a recorded species ranked above an absent species, with half credit for ties (Pearce & Ferrier, 2000). An AUC of 0.5 indicates chance and 1 perfect discrimination. AUC assesses species rankings, rather than occurrence probabilities or the binary counting rule used to reconstruct richness.

Overall AUC was averaged within plots and then equally across all 150 plots. LUI summaries first averaged censuses within plots and brackets, then equally across contributing plots (Fig. S9). We used 9,999 matched whole-plot bootstrap resamples within regions for all rankings, keeping training fits and folds fixed. Bars and ribbons show pointwise 95% percentile intervals. The LUI curves are descriptive, as we did not fit or test an AUC response to LUI.

Ten paired contrasts compared each of the four feasibility rankings with chance and the frequency benchmark, and compared the two interaction scenarios within each combination size. Bonferroni-adjusted 99.5% percentile intervals provided approximate 95% familywise coverage for these contrasts.

### Supplementary figures and tables

#### Figures

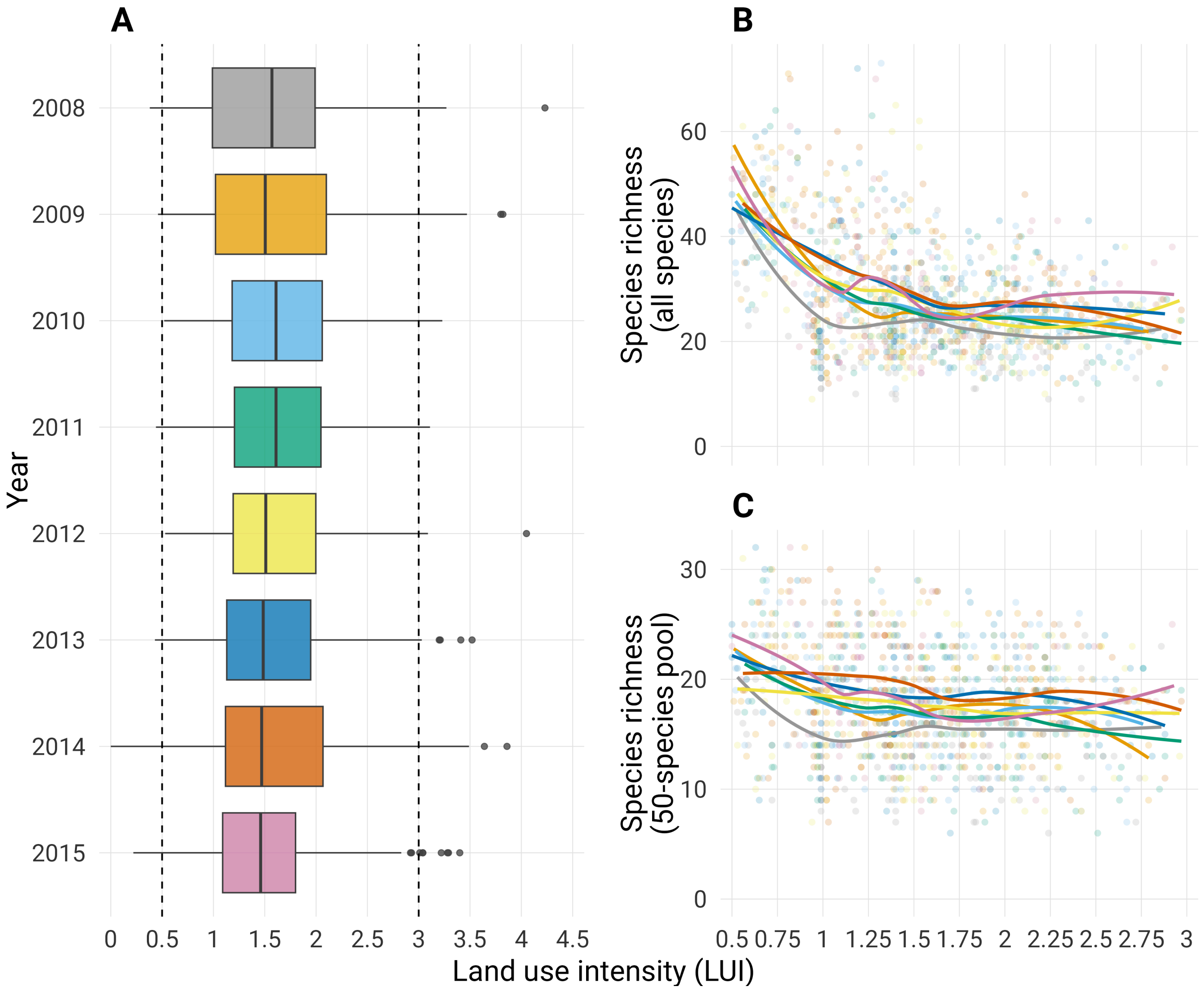


**Figure S1.** Land use intensity (LUI) and observed species richness in 150 grassland plots, 2008–2015. (A) Annual distribution of LUI; dashed lines delimit the analysed range (0.5–3). (B, C) Richness across LUI for all recorded species (B) and the 50-species analysis pool (C). Points represent plot-year censuses; curves are year-specific LOESS fits. Colours identify years and match the boxplots in panel A.

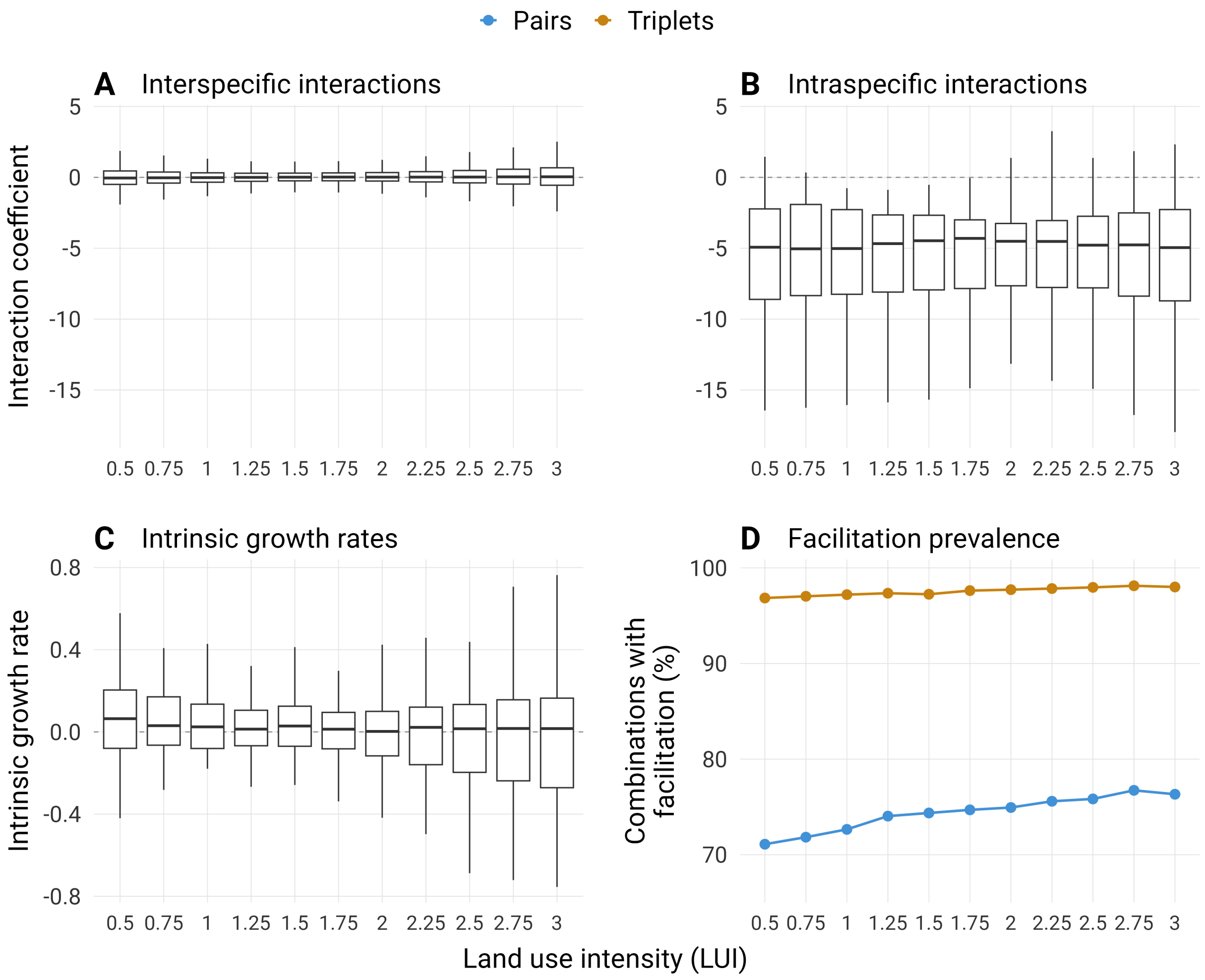


**Figure S2.** Fitted population parameters and facilitation across land use intensity (LUI) for the 50-species pool. (A, B) Fitted interaction coefficients ($b_{ij}$) for the 2,450 directed interspecific interactions (A) and 50 intraspecific interactions (B), shown on the same scale. Negative values indicate competitive effects and positive values facilitative effects. (C) Estimated intrinsic growth rates ($b_{i0}$). Boxes show medians and interquartile ranges, with whiskers extending to the most extreme values within 1.5 interquartile ranges of the box. Outlier points are not shown. (D) Facilitation prevalence: points and lines show the percentage of all 1,225 pairs (blue) and 19,600 triplets (orange) containing at least one facilitative interspecific effect. A single positive directed effect is sufficient to count a combination. Endpoint contrasts and joint tests are reported in Table S3.

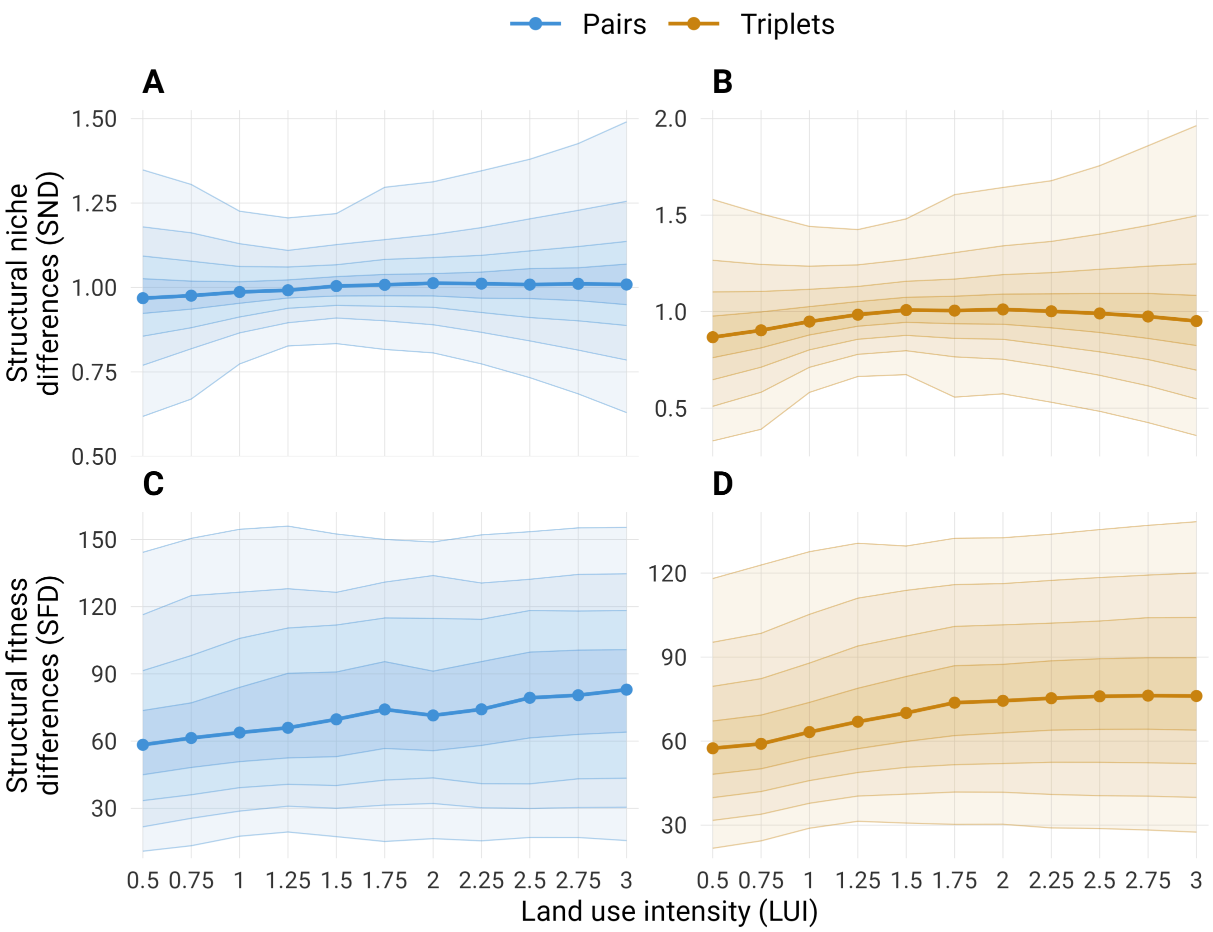


**Figure S3.** Distributions of structural niche differences (SND; A, B) and structural fitness differences (SFD; C, D) across land use intensity (LUI). Pairs are blue and triplets orange. Points and thick lines show medians across all 1,225 pairs or 19,600 triplets from the 50-species pool. Nested bands span the 10th–90th, 20th–80th, 30th–70th and 40th–60th percentiles; thin lines mark their boundaries. Bands describe variation among combinations. These distributions accompany Fig. 3A–B.

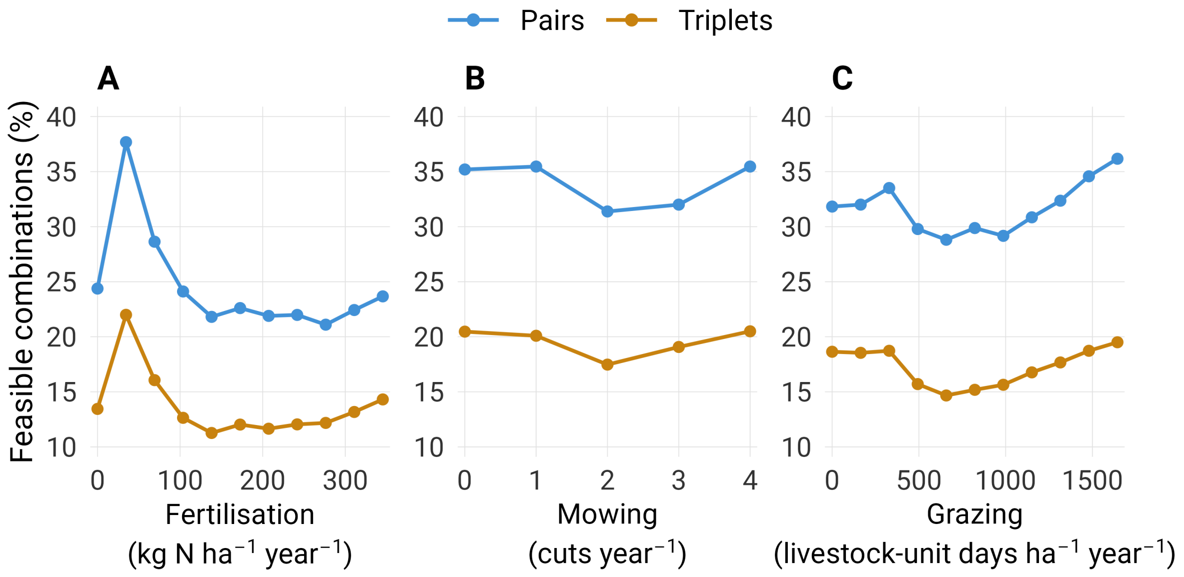


**Figure S4.** Feasibility across fertilisation (A), mowing (B) and grazing (C) for pairs (blue) and triplets (orange). Points show the percentage of feasible combinations from a common pool of 48 species (1,128 pairs and 17,296 triplets), comprising the main pool except *Brachypodium pinnatum* and *Thymus pulegioides*. Components were fitted separately to 1,135 transitions from 150 plots. These associations do not separate the independent effects of correlated management practices. Endpoint contrasts are reported in Table S10 correlations among components in Table S1.

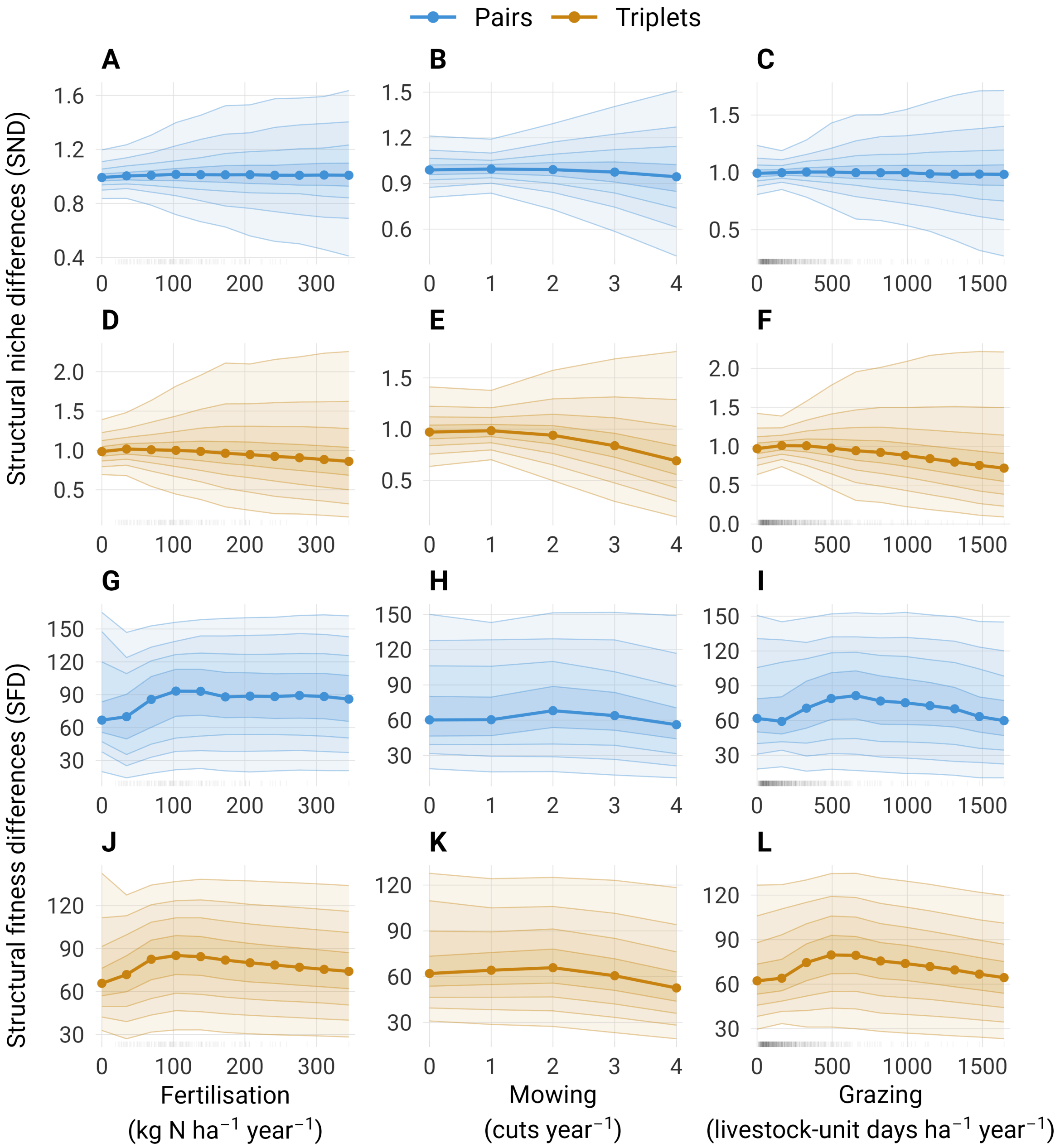


**Figure S5.** Structural niche differences (SND; A–F) and structural fitness differences (SFD; G–L) across fertilisation, mowing and grazing (columns). Rows A–C and G–I show pairs; D–F and J–L show triplets. Points and thick lines show medians; nested bands span the 10th–90th, 20th–80th, 30th–70th and 40th–60th percentiles. Rugs show management values in the population-model data. Analyses use the common 48-species pool defined in Fig. S4. Endpoint median summaries are reported in Table S11 correlations among components in Table S1.

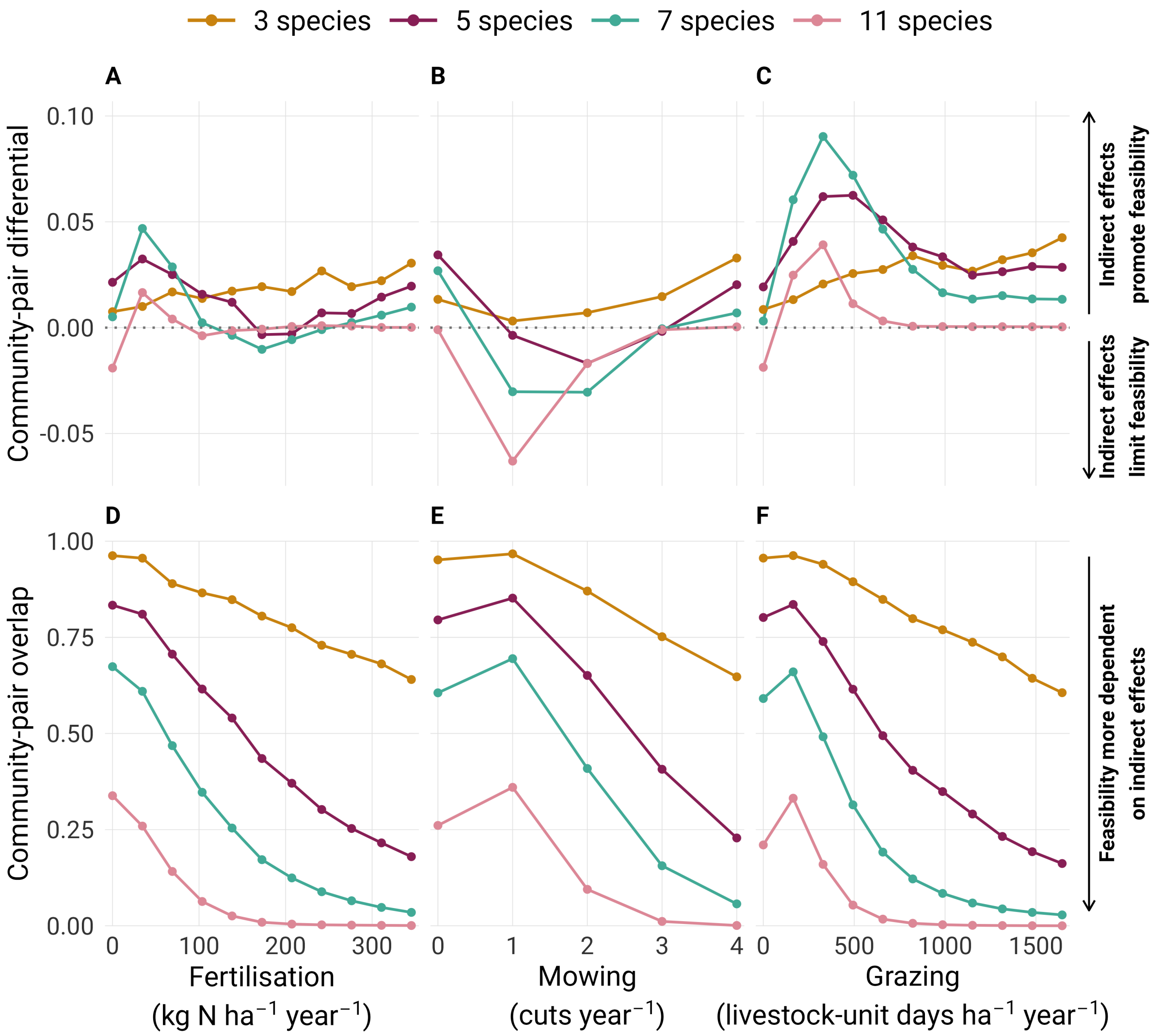


**Figure S6.** Community-pair differential (A–C) and community-pair overlap (D–F) across fertilisation, mowing and grazing (columns), for assemblages of 3, 5, 7 and 11 species from the common 48-species pool. Points and lines show means for a sample of 1,000 assemblages per size. The same combinations are evaluated across levels and components. Positive differential indicates greater full-community feasibility than simultaneous pairwise feasibility; lower overlap indicates greater dependence on indirect effects. Overlap means exclude empty full-community domains. Component-level and assemblage-size contrasts are reported in Table S12 and Table S13.

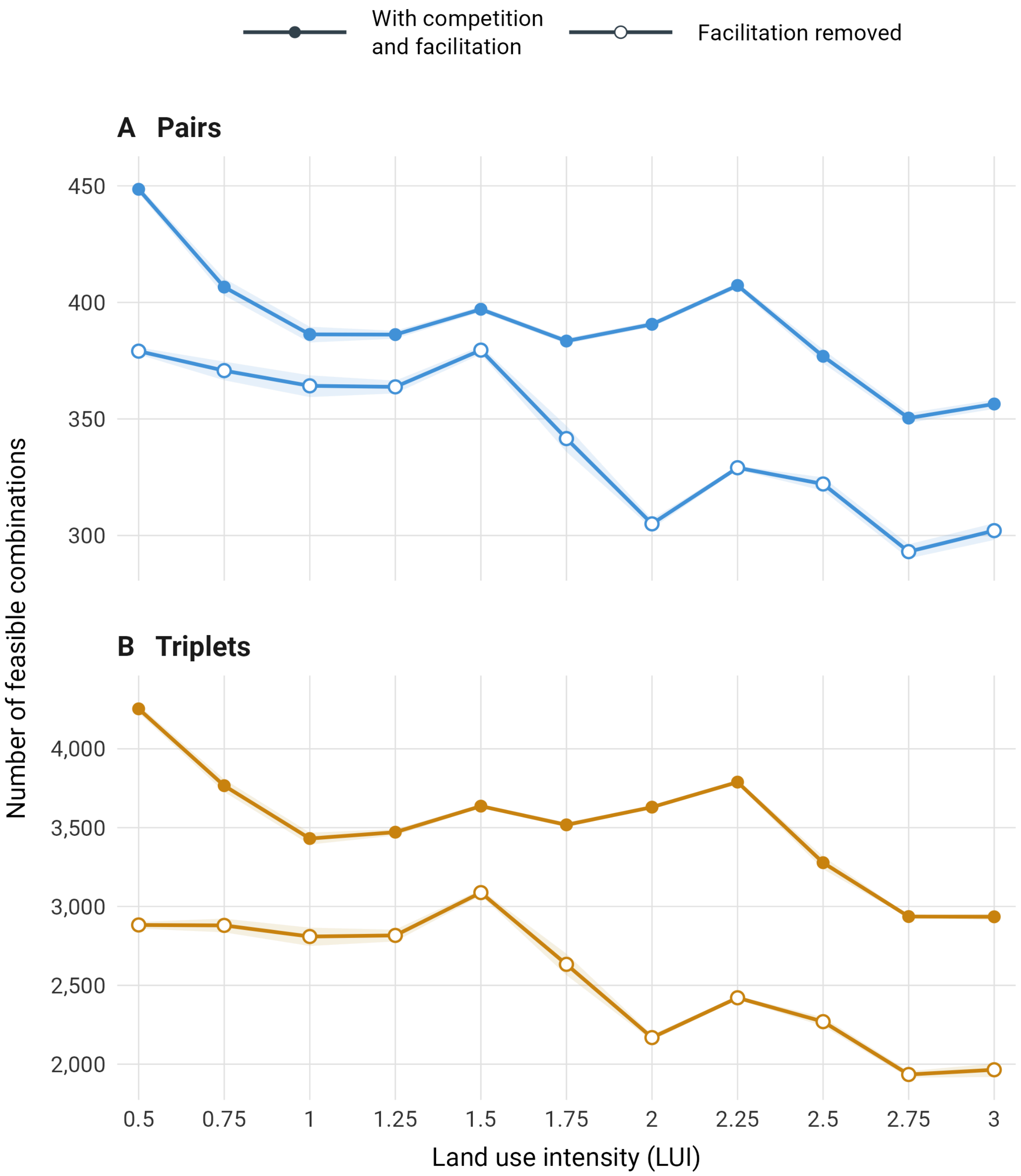


**Figure S7.** Numbers of feasible pairs (A, blue) and triplets (B, orange) across land use intensity (LUI), with competition and facilitation (filled circles) or with facilitation removed (open circles). Both scenarios evaluate the same 50-species pool at each census's actual LUI. Points show means across censuses within plots and quarter-unit LUI brackets, then equally across plots. Shading shows pointwise 95% intervals from 9,999 whole-plot bootstrap resamples within regions, with fitted coefficients fixed. Panels use separate y-axis scales. These counts describe combinations, not distinct species, and accompany the reconstructions in Fig. 5.

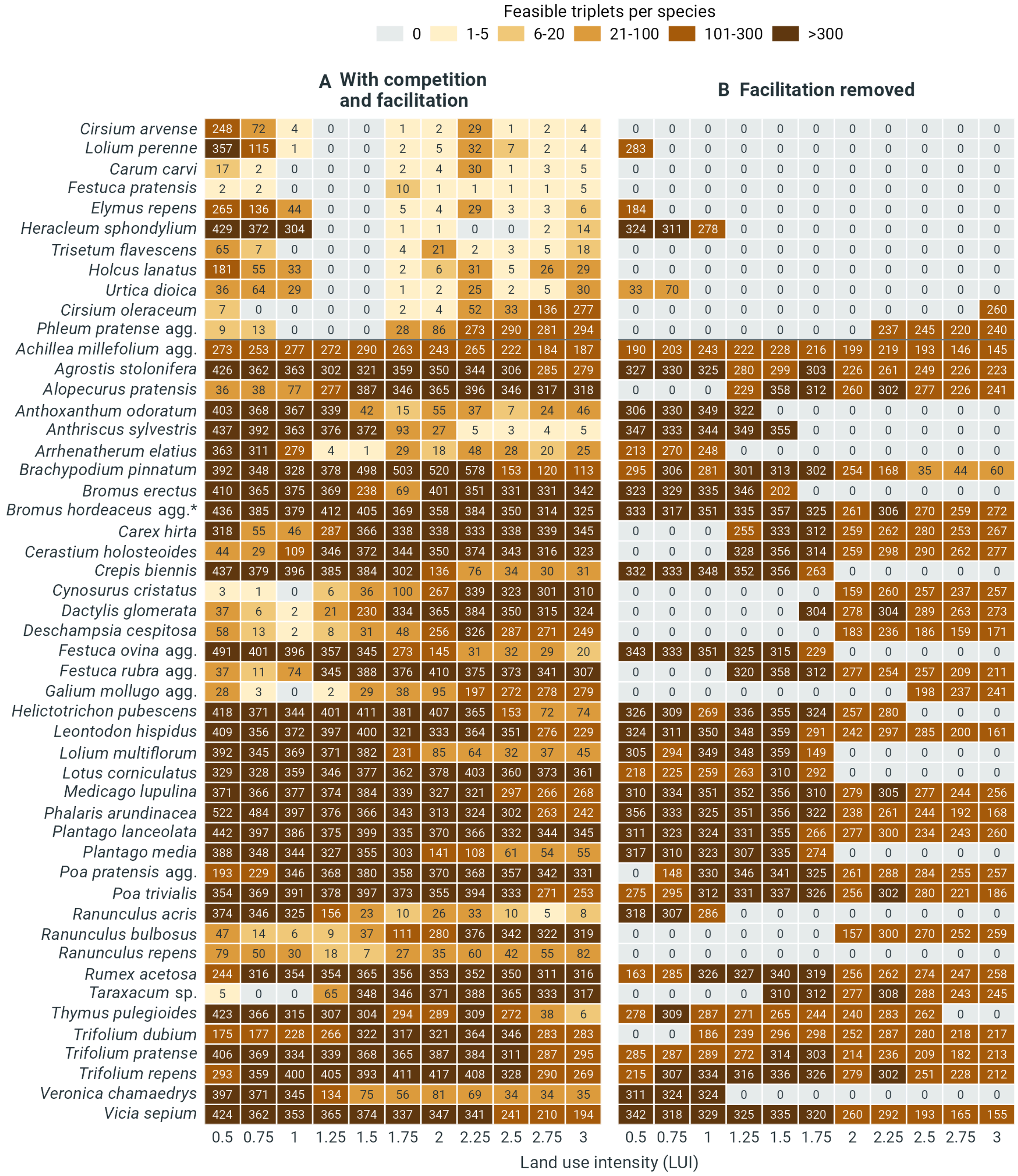


**Figure S8.** Participation of the 50 species in feasible triplets across land use intensity (LUI), with competition and facilitation (A) or with facilitation removed (B). Cells give the number of feasible triplets containing each species; both panels share species order and colour categories. Species above the line account for the rebound between LUI 1.5 and 1.75 with competition and facilitation. Any positive count includes a species in the reconstruction. Counts are evaluated at exact LUI values and do not indicate abundance. *Including *Bromus commutatus*.

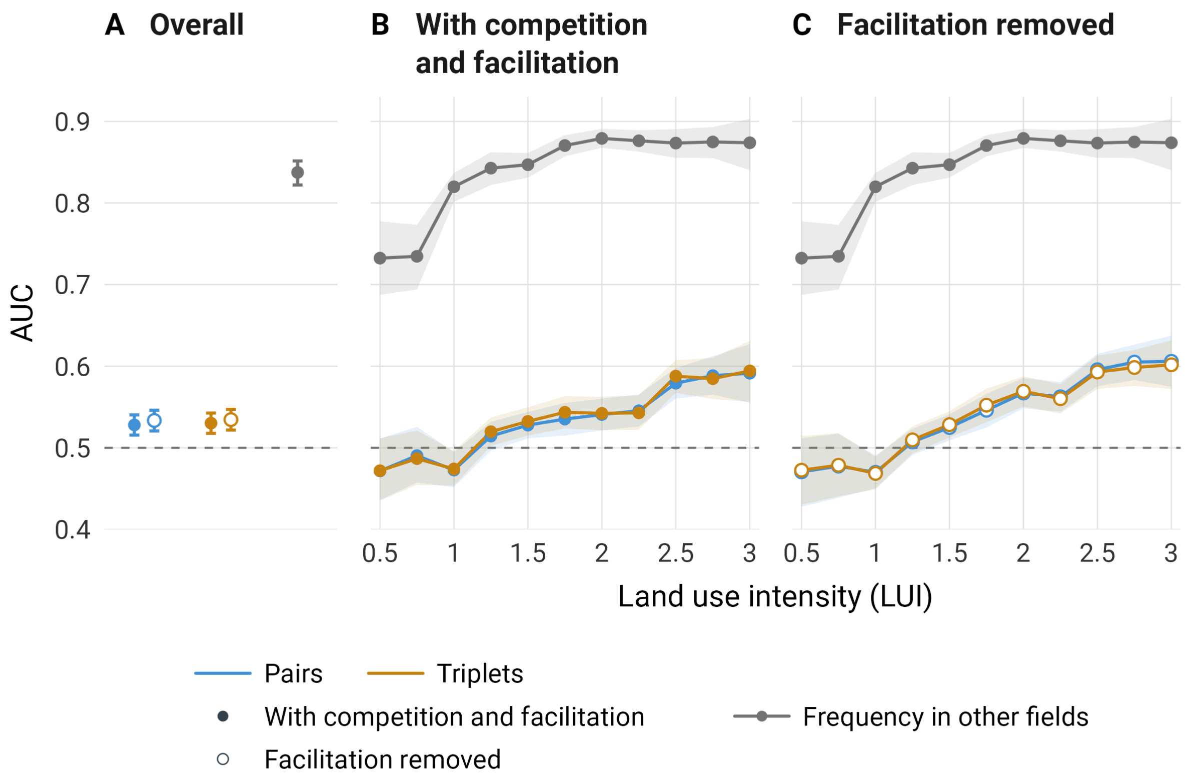


**Figure S9.** Discrimination between recorded and absent species in held-out field censuses, summarised overall (A) and across land use intensity (LUI) with competition and facilitation (B) or with facilitation removed (C). Blue and orange show rankings by participation in feasible pairs and triplets; grey shows the regional training-frequency benchmark. Filled and open circles distinguish the original fitted interactions from facilitation removal; the frequency benchmark is unchanged between scenarios. Both scenarios use the same 50-species pool and training-fitted intrinsic growth rates. Facilitation removal replaces every positive fitted cover effect, including positive intraspecific effects, with $-10^{-6}$ at each census's actual LUI. All records from each test field were withheld during population-model fitting in fivefold cross-validation. AUC (area under the receiver operating characteristic curve) measures how often recorded species receive higher scores than absent species, counting ties as half. An AUC of 0.5 represents chance and 1 perfect discrimination; dashed lines mark 0.5. Overall estimates average censuses within fields and then weight all 150 fields equally. LUI curves first average censuses within fields and quarter-unit LUI brackets, then weight contributing fields equally; points are placed at bracket centres. Bars and ribbons show pointwise 95% intervals from the same 9,999 whole-field bootstrap resamples within regions, with fitted models and folds fixed. The curves describe variation among LUI brackets rather than a fitted LUI effect. Contributing fields vary among brackets, from 21 in the lowest bracket to 14 in the highest.

#### Tables

**Table S1.** Correlations among management components in the common dataset of 1,135 annual transitions from 150 plots. All: pooled regions; A: Schwäbische Alb; H: Hainich-Dün; S: Schorfheide-Chorin. (A) Pearson's $r$ measures linear association. (B) Spearman's $\rho$ measures rank association, with average ranks for ties. Each transition contributes equally to the correlation. Pointwise 95% percentile confidence intervals (CI) use 1,999 bootstrap draws resampling whole plots within regions and retaining every transition of each selected plot. Fisher SE is the bootstrap standard error of the Fisher-transformed correlation, $\operatorname{atanh}(r)$ or $\operatorname{atanh}(\rho)$. Approximate two-sided Wald tests divide the transformed estimate by this SE to assess zero correlation. Reported $p$ values are Benjamini–Hochberg-adjusted across all 24 tests. Pooled correlations combine within-region and between-region variation; regional correlations combine within-plot and between-plot variation.

A) Pearson correlations

| Region | Management pair | Plots | Transitions | Correlation (95% CI) | Fisher SE | *z* | *p* |
|---|---|---|---|---|---|---|---|
| All | Fertilisation–mowing | 150 | 1135 | 0.497 (0.415, 0.575) | 0.055 | 9.853 | <0.001 |
| All | Fertilisation–grazing | 150 | 1135 | -0.165 (-0.264, -0.060) | 0.055 | -3.044 | 0.003 |
| All | Mowing–grazing | 150 | 1135 | -0.480 (-0.545, -0.429) | 0.040 | -13.077 | <0.001 |
| A | Fertilisation–mowing | 50 | 342 | 0.618 (0.478, 0.737) | 0.108 | 6.667 | <0.001 |
| A | Fertilisation–grazing | 50 | 342 | -0.103 (-0.421, 0.149) | 0.170 | -0.611 | 0.542 |
| A | Mowing–grazing | 50 | 342 | -0.448 (-0.617, -0.348) | 0.092 | -5.245 | <0.001 |
| H | Fertilisation–mowing | 50 | 397 | 0.513 (0.366, 0.626) | 0.091 | 6.259 | <0.001 |
| H | Fertilisation–grazing | 50 | 397 | -0.149 (-0.278, -0.075) | 0.056 | -2.652 | 0.010 |
| H | Mowing–grazing | 50 | 397 | -0.308 (-0.421, -0.234) | 0.054 | -5.851 | <0.001 |
| S | Fertilisation–mowing | 50 | 396 | 0.233 (0.044, 0.410) | 0.102 | 2.330 | 0.023 |
| S | Fertilisation–grazing | 50 | 396 | -0.132 (-0.242, -0.014) | 0.059 | -2.260 | 0.026 |
| S | Mowing–grazing | 50 | 396 | -0.658 (-0.711, -0.605) | 0.048 | -16.369 | <0.001 |

B) Spearman rank correlations

| Region | Management pair | Plots | Transitions | Correlation (95% CI) | Fisher SE | *z* | *p* |
|---|---|---|---|---|---|---|---|
| All | Fertilisation–mowing | 150 | 1135 | 0.527 (0.440, 0.609) | 0.061 | 9.678 | <0.001 |
| All | Fertilisation–grazing | 150 | 1135 | -0.276 (-0.375, -0.165) | 0.059 | -4.786 | <0.001 |
| All | Mowing–grazing | 150 | 1135 | -0.684 (-0.735, -0.620) | 0.054 | -15.343 | <0.001 |
| A | Fertilisation–mowing | 50 | 342 | 0.677 (0.525, 0.801) | 0.134 | 6.142 | <0.001 |
| A | Fertilisation–grazing | 50 | 342 | -0.393 (-0.604, -0.163) | 0.135 | -3.085 | 0.003 |
| A | Mowing–grazing | 50 | 342 | -0.713 (-0.817, -0.557) | 0.133 | -6.705 | <0.001 |
| H | Fertilisation–mowing | 50 | 397 | 0.541 (0.387, 0.662) | 0.099 | 6.088 | <0.001 |
| H | Fertilisation–grazing | 50 | 397 | -0.274 (-0.410, -0.120) | 0.081 | -3.498 | <0.001 |
| H | Mowing–grazing | 50 | 397 | -0.482 (-0.586, -0.335) | 0.083 | -6.336 | <0.001 |
| S | Fertilisation–mowing | 50 | 396 | 0.245 (0.043, 0.423) | 0.107 | 2.333 | 0.023 |
| S | Fertilisation–grazing | 50 | 396 | -0.134 (-0.289, 0.041) | 0.088 | -1.530 | 0.131 |
| S | Mowing–grazing | 50 | 396 | -0.804 (-0.845, -0.743) | 0.071 | -15.620 | <0.001 |

**Table S2.** Occurrence frequency (%) of the 50 analysis taxa, ranked by overall frequency. Values are percentages of 1,191 complete plot-year censuses (150 plots, 2008–2015) with positive cover. Regions: A, Schwäbische Alb (*n* = 396); H, Hainich-Dün (*n* = 397); S, Schorfheide-Chorin (*n* = 398). LUI tertiles: low, ≤1.31 (*n* = 384); mid, >1.31–1.83 (*n* = 379); high, >1.83 (*n* = 378). Fifty censuses lacking LUI contribute only to overall and regional frequencies. agg., species aggregate; sp., genus-level identification.

| Scientific name | Overall | A | H | S | Low LUI | Mid LUI | High LUI |
|---|---|---|---|---|---|---|---|
| *Taraxacum* sp. | 91.7 | 86.1 | 96.0 | 93.0 | 83.9 | 94.5 | 97.6 |
| *Poa pratensis* agg. | 86.3 | 73.0 | 94.0 | 92.0 | 86.7 | 88.1 | 88.6 |
| *Dactylis glomerata* | 83.3 | 85.6 | 91.4 | 72.9 | 81.2 | 81.5 | 86.8 |
| *Cerastium holosteoides* | 79.8 | 78.5 | 79.6 | 81.4 | 71.1 | 85.8 | 83.3 |
| *Trifolium repens* | 77.5 | 86.9 | 81.6 | 64.1 | 70.1 | 77.0 | 83.3 |
| *Plantago lanceolata* | 68.6 | 80.6 | 75.8 | 49.5 | 71.1 | 65.4 | 67.2 |
| *Poa trivialis* | 67.9 | 72.2 | 54.9 | 76.6 | 54.9 | 72.6 | 74.9 |
| *Trifolium pratense* | 61.7 | 86.6 | 69.3 | 29.4 | 61.2 | 59.9 | 60.1 |
| *Achillea millefolium* agg. | 54.4 | 47.5 | 68.3 | 47.5 | 64.8 | 53.8 | 45.2 |
| *Festuca pratensis* | 54.4 | 52.8 | 83.9 | 26.6 | 60.7 | 48.8 | 51.9 |
| *Lolium perenne* | 54.4 | 60.4 | 52.1 | 50.8 | 32.6 | 59.9 | 69.8 |
| *Veronica chamaedrys* | 51.7 | 65.2 | 56.4 | 33.7 | 55.2 | 50.9 | 47.1 |
| *Galium mollugo* agg. | 47.0 | 68.2 | 60.2 | 12.8 | 48.2 | 43.5 | 45.5 |
| *Alopecurus pratensis* | 46.8 | 53.8 | 53.4 | 33.2 | 31.8 | 47.5 | 59.8 |
| *Elymus repens* | 44.2 | 0.3 | 57.9 | 74.4 | 44.8 | 49.3 | 44.4 |
| *Bromus hordeaceus* agg.* | 41.3 | 31.8 | 48.1 | 44.0 | 29.4 | 47.2 | 50.0 |
| *Trisetum flavescens* | 41.0 | 79.5 | 41.6 | 2.0 | 41.4 | 33.8 | 42.6 |
| *Arrhenatherum elatius* | 40.4 | 37.4 | 54.2 | 29.6 | 49.7 | 37.5 | 33.3 |
| *Festuca rubra* agg. | 39.9 | 44.7 | 47.9 | 27.1 | 53.4 | 35.6 | 28.8 |
| *Ranunculus repens* | 38.4 | 18.9 | 34.8 | 61.3 | 39.1 | 43.8 | 34.4 |
| *Ranunculus acris* | 37.9 | 80.1 | 22.2 | 11.6 | 33.6 | 35.6 | 38.4 |
| *Anthriscus sylvestris* | 37.7 | 58.8 | 37.0 | 17.3 | 29.2 | 36.1 | 44.7 |
| *Heracleum sphondylium* | 35.2 | 54.3 | 19.9 | 31.4 | 21.6 | 35.9 | 46.0 |
| *Ranunculus bulbosus* | 34.0 | 37.6 | 39.5 | 24.9 | 50.5 | 30.1 | 20.9 |
| *Rumex acetosa* | 33.0 | 68.9 | 11.6 | 18.6 | 21.9 | 33.5 | 38.1 |
| *Phleum pratense* agg. | 31.9 | 23.5 | 47.4 | 24.9 | 22.9 | 33.2 | 39.2 |
| *Crepis biennis* | 29.8 | 29.0 | 59.9 | 0.5 | 26.8 | 28.2 | 34.4 |
| *Vicia sepium* | 28.9 | 65.2 | 17.4 | 4.3 | 24.5 | 27.2 | 29.6 |
| *Holcus lanatus* | 27.0 | 17.4 | 15.6 | 47.7 | 35.9 | 27.4 | 17.2 |
| *Cirsium arvense* | 24.6 | 3.0 | 26.7 | 44.0 | 27.1 | 29.6 | 20.4 |
| *Lotus corniculatus* | 24.4 | 32.1 | 37.3 | 4.0 | 44.3 | 17.4 | 9.5 |
| *Medicago lupulina* | 23.9 | 23.7 | 39.0 | 9.0 | 38.5 | 18.7 | 14.0 |
| *Plantago media* | 23.3 | 46.0 | 23.4 | 0.5 | 41.4 | 15.0 | 10.3 |
| *Trifolium dubium* | 20.5 | 22.0 | 20.4 | 19.1 | 26.8 | 21.6 | 12.4 |
| *Carum carvi* | 19.7 | 34.3 | 24.9 | 0.0 | 18.0 | 14.0 | 25.4 |
| *Helictotrichon pubescens* | 19.6 | 39.6 | 17.9 | 1.5 | 28.6 | 16.1 | 10.6 |
| *Cynosurus cristatus* | 18.6 | 41.2 | 10.3 | 4.3 | 25.0 | 18.5 | 8.2 |
| *Anthoxanthum odoratum* | 17.7 | 41.7 | 9.8 | 1.8 | 26.3 | 15.8 | 6.9 |
| *Festuca ovina* agg. | 14.2 | 18.2 | 24.2 | 0.3 | 32.8 | 6.3 | 2.6 |
| *Deschampsia cespitosa* | 12.0 | 2.3 | 12.6 | 21.1 | 20.1 | 8.7 | 8.5 |
| *Agrostis stolonifera* | 11.9 | 0.3 | 14.4 | 21.1 | 15.9 | 11.1 | 10.3 |
| *Bromus erectus* | 11.9 | 23.7 | 12.1 | 0.0 | 24.5 | 6.6 | 2.6 |
| *Carex hirta* | 11.8 | 0.3 | 0.0 | 35.2 | 16.7 | 12.4 | 7.9 |
| *Urtica dioica* | 11.6 | 3.8 | 4.0 | 26.9 | 14.3 | 15.0 | 6.6 |
| *Leontodon hispidus* | 10.7 | 12.9 | 19.1 | 0.0 | 25.0 | 5.3 | 1.9 |
| *Cirsium oleraceum* | 9.8 | 0.0 | 2.3 | 27.1 | 13.5 | 11.6 | 5.6 |
| *Thymus pulegioides* | 9.7 | 23.5 | 5.8 | 0.0 | 21.9 | 5.3 | 0.0 |
| *Brachypodium pinnatum* | 8.7 | 14.4 | 11.8 | 0.0 | 20.6 | 4.0 | 0.8 |
| *Lolium multiflorum* | 7.5 | 0.0 | 10.8 | 11.6 | 5.5 | 8.7 | 9.3 |
| *Phalaris arundinacea* | 7.0 | 0.0 | 0.0 | 20.9 | 8.3 | 9.0 | 4.5 |

*Including *Bromus commutatus*

**Table S3.** Changes in fitted interspecific effects, intraspecific effects, intrinsic growth rates and facilitation prevalence from LUI 0.5 to 3 for the 50-species pool in Fig. S2. (A) Changes are high minus low; negative interaction-effect changes indicate a shift towards greater growth suppression, and negative growth-rate changes indicate lower intrinsic growth. Positive-effect prevalence is the percentage of all 1,225 pairs or 19,600 triplets containing at least one positive directed interspecific effect; its changes are in percentage points (pp). Confidence intervals (CI) are pointwise 95% bootstrap percentile intervals, and SE is the bootstrap standard error, from 998 successful joint parametric bootstrap draws out of 999. Simulations preserve the fitted region/plot, temporal and working cross-species covariance, and refit the population models. Approximate two-sided Wald tests use $z$ = change/SE. (B) Joint Wald tests use the bootstrap covariance and test whether both endpoint changes are zero; df denotes degrees of freedom. Benjamini–Hochberg adjustment is applied separately to the eight tests in A and the two tests in B.

A) Endpoint changes

| **Quantity** | **Change (95% CI)** | **Bootstrap SE** | ***z*** | ***p*** |
|---|---|---|---|---|
| Mean interspecific coefficient | 0.028 (-0.208, 0.274) | 0.123 | 0.227 | 0.938 |
| Median interspecific coefficient | 0.087 (-0.076, 0.282) | 0.092 | 0.940 | 0.556 |
| Mean intraspecific coefficient | -1.038 (-1.699, -0.390) | 0.337 | -3.080 | 0.017 |
| Median intraspecific coefficient | -0.026 (-0.818, 0.907) | 0.456 | -0.056 | 0.955 |
| Mean intrinsic growth rate | -0.106 (-0.278, 0.070) | 0.091 | -1.164 | 0.489 |
| Median intrinsic growth rate | -0.048 (-0.294, 0.090) | 0.097 | -0.498 | 0.824 |
| Positive effects: pairs (pp) | 5.224 (-3.353, 10.373) | 3.396 | 1.539 | 0.356 |
| Positive effects: triplets (pp) | 1.153 (-0.817, 2.327) | 0.768 | 1.501 | 0.356 |

B) Joint endpoint tests

| **Quantities** | **Wald $\chi^2$** | **df** | ***p*** |
|---|---|---|---|
| Mean inter- and intraspecific coefficients | 10.849 | 2 | 0.009 |
| Positive effects in pairs and triplets | 2.373 | 2 | 0.305 |

**Table S4.** Feasibility at LUI 0.5 and 3 and its net change in Fig. 2A, across all 1,225 pairs and 19,600 triplets from the 50-species pool. Change is feasibility at LUI 3 minus feasibility at LUI 0.5; changes, standard errors (SE) and confidence intervals (CI) are in percentage points. Parentheses give pointwise 95% percentile intervals from 998 successful joint parametric bootstrap draws out of 999 attempts, propagating population-model uncertainty. Approximate two-sided Wald tests use $z = \mathtt{change} / \mathtt{SE}$ with a standard normal reference and test zero net change. Tests concern the endpoint difference, not the shape of the LUI response. Species population models (Equation 1) used Gaussian REML, random intercepts for region and plots within regions, and within-plot AR(1) errors. The bootstrap refits these models; the contrasts are derived from their coefficients.

| **Assemblage** | **LUI 0.5 (%)** | **LUI 3 (%)** | **Change (95% CI)** | **SE** | ***z*** | ***p*** |
|---|---|---|---|---|---|---|
| Pairs | 36.816 | 29.143 | -7.673 (-25.965, 9.224) | 8.918 | -0.860 | 0.390 |
| Triplets | 21.990 | 15.036 | -6.954 (-18.503, 6.172) | 6.391 | -1.088 | 0.277 |

**Table S5.** Contrasts in median SND and SFD for Fig. 3A–B. Low, middle and high denote LUI 0.5, 1.75 and 3. Fitted changes compare empirical medians across all pairs or triplets of the 50-species pool; bias-corrected changes subtract the estimated bootstrap bias. Bootstrap standard errors (SE) use 998 successful joint parametric draws, refitting the species population models described in Table S4. Pointwise 95% intervals use critical values from 750 calibration datasets, with 199 inner refits attempted per dataset; 250 separately generated datasets assess coverage. For testing zero change, $T$ is the absolute bias-corrected change divided by SE. Approximate two-sided $p$ values use $(1 + b)/751$, where $b$ counts calibration scores at least as large as $T$, followed by Holm adjustment across the 12 median comparisons. The unadjusted simulation resolution is 1/751 (approximately 0.0013). Intervals are pointwise, whereas the adjustment applies to $p$ values. Intervals and tests retain the assumptions of the fitted-model calibration.

| **Metric** | **Assemblage** | **Contrast** | **Fitted change** | **Bias-corrected change** | **Bootstrap SE** | ***T*** | **95% interval** | ***p*** |
|---|---|---|---|---|---|---|---|---|
| SND | Pairs | High - low | 0.040 | 0.052 | 0.034 | 1.543 | -0.025 to 0.129 | 1.000 |
| SND | Pairs | Middle - low | 0.040 | 0.048 | 0.020 | 2.407 | -<0.001 to 0.096 | 0.655 |
| SND | Pairs | Middle - high | -<0.001 | -0.004 | 0.017 | 0.241 | -0.048 to 0.039 | 1.000 |
| SND | Triplets | High - low | 0.084 | 0.132 | 0.084 | 1.558 | -0.089 to 0.353 | 1.000 |
| SND | Triplets | Middle - low | 0.138 | 0.094 | 0.053 | 1.779 | -0.032 to 0.221 | 1.000 |
| SND | Triplets | Middle - high | 0.054 | -0.037 | 0.042 | 0.887 | -0.182 to 0.107 | 1.000 |
| SFD | Pairs | High - low | 24.612 | 30.515 | 17.857 | 1.709 | -28.550 to 89.581 | 1.000 |
| SFD | Pairs | Middle - low | 15.777 | 15.475 | 14.201 | 1.090 | -29.139 to 60.088 | 1.000 |
| SFD | Pairs | Middle - high | -8.834 | -15.041 | 9.735 | 1.545 | -49.183 to 19.101 | 1.000 |
| SFD | Triplets | High - low | 18.674 | 23.779 | 12.774 | 1.862 | -18.613 to 66.171 | 1.000 |
| SFD | Triplets | Middle - low | 16.285 | 16.384 | 10.146 | 1.615 | -13.476 to 46.245 | 1.000 |
| SFD | Triplets | Middle - high | -2.389 | -7.395 | 6.464 | 1.144 | -27.029 to 12.239 | 1.000 |

**Table S6.** Observed-minus-unobserved feasibility in Fig. 2B, in percentage points. Each census compares entirely observed combinations with combinations containing exactly one unobserved species and otherwise observed partners; positive values indicate higher feasibility for observed-only combinations. Separate Gaussian REML models for pairs and triplets use 1,118 censuses from 150 plots: difference ~ (LUI −1.75) + region + year, with categorical region and year, a plot random intercept, temporal AR(1) correlation and region-specific residual variances. (A) Contrasts average regions and years equally; low and high denote LUI 0.5 and 3. Parentheses give pointwise 95% *t* confidence intervals; SE: standard error; df: containment degrees of freedom. Two-sided *t* tests use Holm adjustment across the six contrasts here and twelve in Table S7. (B) Intercepts and LUI slopes are reported with SE in parentheses; SD denotes standard deviation. Intercepts refer to LUI 1.75, region A and year 2008, with plot effect zero. Slopes are changes per unit LUI. A: Schwäbische Alb; H: Hainich-Dün; S: Schorfheide-Chorin. AR(1) is the residual correlation between consecutive years within a plot.

A) Contrasts

| **Assemblage** | **Contrast** | **Estimate (95% CI)** | **SE** | **df** | ***t*** | ***p*** |
|---|---|---|---|---|---|---|
| Pairs | At low LUI | -4.767 (-6.569, -2.965) | 0.912 | 147 | -5.228 | <0.001 |
| Pairs | At high LUI | 11.340 (9.326, 13.354) | 1.019 | 147 | 11.128 | <0.001 |
| Pairs | High - low | 16.107 (13.603, 18.611) | 1.276 | 960 | 12.623 | <0.001 |
| Triplets | At low LUI | -1.787 (-3.036, -0.538) | 0.632 | 147 | -2.828 | 0.005 |
| Triplets | At high LUI | 6.395 (4.996, 7.794) | 0.708 | 147 | 9.032 | <0.001 |
| Triplets | High - low | 8.182 (6.425, 9.939) | 0.895 | 960 | 9.138 | <0.001 |

B) Model summary

| **Parameter** | **Pairs** | **Triplets** |
|---|---|---|
| Intercept (SE) | 4.983 (1.328) | 3.311 (0.917) |
| LUI slope (SE) | 6.443 (0.510) | 3.273 (0.358) |
| Plot SD | 8.401 | 5.730 |
| Residual SD: A | 5.722 | 4.095 |
| Residual SD: H | 6.426 | 4.627 |
| Residual SD: S | 5.829 | 4.133 |
| AR(1) correlation | 0.261 | 0.301 |

**Table S7.** Observed-minus-unobserved SND and SFD in Fig. 3C–D. Each census compares mean metrics for entirely observed combinations with combinations containing exactly one unobserved species and otherwise observed partners. Four Gaussian REML models use 1,118 censuses from 150 plots: difference ~ (LUI −1.75) + region + year, with categorical region and year, a plot random intercept, temporal AR(1) correlation and region-specific residual variances. (A) Contrasts average regions and years equally; low and high denote LUI 0.5 and 3. Parentheses give pointwise 95% *t* confidence intervals; SE: standard error; df: containment degrees of freedom. Two-sided *t* tests use Holm adjustment across the 18 contrasts here and in Table S6. (B) Intercepts and LUI slopes are reported with SE in parentheses; SD denotes standard deviation. Intercepts refer to LUI 1.75, region A and year 2008, with plot effect zero. Slopes are changes per unit LUI. A: Schwäbische Alb; H: Hainich-Dün; S: Schorfheide-Chorin. AR(1) is the residual correlation between consecutive years within a plot.

A) Contrasts

| Metric | Assemblage | Contrast | Estimate (95% CI) | SE | df | *t* | *p* |
|---|---|---|---|---|---|---|---|
| SND | Pairs | At low LUI | 0.044 (0.040, 0.048) | 0.002 | 147 | 21.124 | <0.001 |
| SND | Pairs | At high LUI | 0.009 (0.004, 0.014) | 0.002 | 147 | 3.623 | 0.001 |
| SND | Pairs | High - low | -0.035 (-0.042, -0.028) | 0.004 | 960 | -9.767 | <0.001 |
| SND | Triplets | At low LUI | 0.091 (0.082, 0.100) | 0.004 | 147 | 20.397 | <0.001 |
| SND | Triplets | At high LUI | 0.019 (0.009, 0.029) | 0.005 | 147 | 3.602 | 0.001 |
| SND | Triplets | High - low | -0.072 (-0.087, -0.057) | 0.008 | 960 | -9.529 | <0.001 |
| SFD | Pairs | At low LUI | 9.329 (7.531, 11.128) | 0.910 | 147 | 10.253 | <0.001 |
| SFD | Pairs | At high LUI | -14.656 (-16.658, -12.653) | 1.013 | 147 | -14.464 | <0.001 |
| SFD | Pairs | High - low | -23.985 (-26.440, -21.531) | 1.251 | 960 | -19.177 | <0.001 |
| SFD | Triplets | At low LUI | 5.735 (4.649, 6.820) | 0.549 | 147 | 10.443 | <0.001 |
| SFD | Triplets | At high LUI | -9.190 (-10.404, -7.976) | 0.614 | 147 | -14.958 | <0.001 |
| SFD | Triplets | High - low | -14.925 (-16.441, -13.409) | 0.773 | 960 | -19.319 | <0.001 |

B) Model summary

| Parameter | SND: pairs | SND: triplets | SFD: pairs | SFD: triplets |
|---|---|---|---|---|
| Intercept (SE) | 0.0149 (0.0027) | 0.0282 (0.0057) | -5.806 (1.328) | -3.509 (0.789) |
| LUI slope (SE) | -0.0140 (0.0014) | -0.0289 (0.0030) | -9.594 (0.500) | -5.970 (0.309) |
| Plot SD | 0.0136 | 0.0290 | 8.326 | 4.931 |
| Residual SD: A | 0.0161 | 0.0345 | 5.589 | 3.297 |
| Residual SD: H | 0.0202 | 0.0415 | 6.433 | 3.962 |
| Residual SD: S | 0.0224 | 0.0538 | 6.628 | 4.304 |
| AR(1) correlation | 0.364 | 0.406 | 0.388 | 0.383 |

**Table S8.** LUI contrasts in community-pair differential (A) and community-pair overlap (B) for Fig. 4. Low, middle and high denote LUI 0.5, 1.75 and 3. Each estimate averages paired contrasts from ten independently sampled sets of 1,000 assemblages per size from the 50-species pool; each set is held constant across LUI. Monte Carlo standard error (MC SE) is $s/\sqrt{10}$, where $s$ is the SD of the ten contrasts. Parentheses give pointwise 95% Monte Carlo intervals, calculated as mean $\pm t_{0.975,9}$ × MC SE. Agreement is the number of samples with the same sign as the mean contrast. Intervals describe precision from sampling combinations at the fitted coefficients; they do not propagate population-model uncertainty. The two middle contrasts describe the rise and fall of the overlap curve, whereas high minus low measures net change. A positive differential change indicates a shift towards greater promotion of feasibility; a negative overlap change indicates greater dependence on indirect effects. Overlap means exclude empty full feasibility domains separately at each LUI.

A) Community-pair differential

| Species | Contrast | Change (95% MC interval) | MC SE | Agreement |
|---|---|---|---|---|
| 3 | Middle - low | -0.0088 (-0.0119, -0.0058) | 0.0014 | 10/10 |
| 3 | High - middle | 0.0153 (0.0116, 0.0189) | 0.0016 | 10/10 |
| 3 | High - low | 0.0065 (0.0010, 0.0119) | 0.0024 | 8/10 |
| 5 | Middle - low | 0.0035 (-0.0004, 0.0073) | 0.0017 | 8/10 |
| 5 | High - middle | 0.0146 (0.0100, 0.0192) | 0.0020 | 10/10 |
| 5 | High - low | 0.0180 (0.0138, 0.0223) | 0.0019 | 10/10 |
| 7 | Middle - low | 0.0185 (0.0151, 0.0219) | 0.0015 | 10/10 |
| 7 | High - middle | -0.0071 (-0.0113, -0.0030) | 0.0018 | 9/10 |
| 7 | High - low | 0.0113 (0.0074, 0.0153) | 0.0018 | 10/10 |
| 11 | Middle - low | 0.0070 (0.0041, 0.0099) | 0.0013 | 10/10 |
| 11 | High - middle | -0.0057 (-0.0081, -0.0033) | 0.0011 | 10/10 |
| 11 | High - low | 0.0013 (0.0006, 0.0020) | 0.0003 | 9/10 |

B) Community-pair overlap

| Species | Contrast | Change (95% MC interval) | MC SE | Agreement |
|---|---|---|---|---|
| 3 | Middle - low | 0.1199 (0.1150, 0.1249) | 0.0022 | 10/10 |
| 3 | High - middle | -0.1392 (-0.1478, -0.1307) | 0.0038 | 10/10 |
| 3 | High - low | -0.0193 (-0.0274, -0.0111) | 0.0036 | 10/10 |
| 5 | Middle - low | 0.2764 (0.2656, 0.2872) | 0.0048 | 10/10 |
| 5 | High - middle | -0.3126 (-0.3172, -0.3081) | 0.0020 | 10/10 |
| 5 | High - low | -0.0362 (-0.0486, -0.0238) | 0.0055 | 10/10 |
| 7 | Middle - low | 0.3197 (0.3155, 0.3238) | 0.0018 | 10/10 |
| 7 | High - middle | -0.3603 (-0.3679, -0.3526) | 0.0034 | 10/10 |
| 7 | High - low | -0.0406 (-0.0474, -0.0338) | 0.0030 | 10/10 |
| 11 | Middle - low | 0.1901 (0.1862, 0.1939) | 0.0017 | 10/10 |
| 11 | High - middle | -0.2029 (-0.2064, -0.1993) | 0.0016 | 10/10 |
| 11 | High - low | -0.0128 (-0.0157, -0.0100) | 0.0013 | 10/10 |

**Table S9.** Differences in community-pair differential (A) and community-pair overlap (B) between adjacent assemblage sizes in Fig. 4, at low (0.5), middle (1.75) and high (3) LUI. Each size contrast subtracts the smaller-assemblage mean from the larger-assemblage mean (e.g., 5 minus 3 species). Estimates average contrasts from ten independent sampling runs, each containing 1,000 assemblages per size from the 50-species pool. Monte Carlo standard errors (MC SE), pointwise 95% Monte Carlo intervals and sign agreement are calculated as in Table S8. Agreement gives the number of runs with the same sign as the mean contrast. These intervals quantify combination-sampling precision, not population-model uncertainty. Positive differential differences indicate greater promotion of feasibility in larger assemblages; negative overlap differences indicate greater dependence on indirect effects. Overlap means exclude empty full feasibility domains separately at each size and LUI. Assemblages were sampled separately at each size; these comparisons do not represent adding species to a particular assemblage.

A) Community-pair differential

| Size contrast | LUI | Difference (95% MC interval) | MC SE | Agreement |
|---|---|---|---|---|
| 5 - 3 | 0.5 | 0.0059 (0.0014, 0.0103) | 0.0019 | 9/10 |
| 5 - 3 | 1.75 | 0.0182 (0.0144, 0.0219) | 0.0017 | 10/10 |
| 5 - 3 | 3 | 0.0174 (0.0098, 0.0251) | 0.0034 | 10/10 |
| 7 - 5 | 0.5 | -0.0130 (-0.0169, -0.0090) | 0.0017 | 10/10 |
| 7 - 5 | 1.75 | 0.0020 (-0.0025, 0.0066) | 0.0020 | 6/10 |
| 7 - 5 | 3 | -0.0197 (-0.0239, -0.0155) | 0.0018 | 10/10 |
| 11 - 7 | 0.5 | -0.0115 (-0.0143, -0.0087) | 0.0012 | 10/10 |
| 11 - 7 | 1.75 | -0.0229 (-0.0269, -0.0190) | 0.0018 | 10/10 |
| 11 - 7 | 3 | -0.0215 (-0.0240, -0.0190) | 0.0011 | 10/10 |

B) Community-pair overlap

| Size contrast | LUI | Difference (95% MC interval) | MC SE | Agreement |
|---|---|---|---|---|
| 5 - 3 | 0.5 | -0.3327 (-0.3432, -0.3222) | 0.0046 | 10/10 |
| 5 - 3 | 1.75 | -0.1762 (-0.1804, -0.1720) | 0.0019 | 10/10 |
| 5 - 3 | 3 | -0.3496 (-0.3598, -0.3395) | 0.0045 | 10/10 |
| 7 - 5 | 0.5 | -0.2646 (-0.2750, -0.2542) | 0.0046 | 10/10 |
| 7 - 5 | 1.75 | -0.2213 (-0.2274, -0.2153) | 0.0027 | 10/10 |
| 7 - 5 | 3 | -0.2690 (-0.2764, -0.2615) | 0.0033 | 10/10 |
| 11 - 7 | 0.5 | -0.2030 (-0.2083, -0.1977) | 0.0023 | 10/10 |
| 11 - 7 | 1.75 | -0.3326 (-0.3413, -0.3239) | 0.0039 | 10/10 |
| 11 - 7 | 3 | -0.1752 (-0.1823, -0.1682) | 0.0031 | 10/10 |

**Table S10.** Feasibility at low and high management intensity and its net change in Fig. S4, across all 1,128 pairs and 17,296 triplets from the common 48-species pool defined in that figure. Low is zero for each component; high is 345.35 kg N $ha^{-1}$ $year^{-1}$ for fertilisation, 4 cuts $year^{-1}$ for mowing and 1,644.167 livestock-unit days $ha^{-1}$ $year^{-1}$ for grazing. Change is high minus low; changes, standard errors (SE) and confidence intervals (CI) are in percentage points. Parentheses give pointwise 95% percentile intervals from 999 successful joint parametric bootstrap draws per component, propagating population-model uncertainty. Approximate two-sided Wald tests use $z$ = change / SE with a standard normal reference to test zero endpoint change. Each component replaces LUI in Equation 1, with the Gaussian REML and covariance structure described in Table S4. Models were fitted separately using the same 1,135 transitions from 150 plots. These tests assess net endpoint differences, not curve shapes or the independent effects of correlated management practices.

| Component | Assemblage | Low (%) | High (%) | Change (95% CI) | SE | *z* | *p* |
|---|---|---|---|---|---|---|---|
| Fertilisation | Pairs | 24.379 | 23.670 | -0.709 (-17.110, 12.416) | 7.568 | -0.094 | 0.925 |
| Fertilisation | Triplets | 13.448 | 14.310 | 0.861 (-12.818, 8.793) | 5.582 | 0.154 | 0.877 |
| Mowing | Pairs | 35.195 | 35.461 | 0.266 (-13.573, 18.887) | 8.440 | 0.032 | 0.975 |
| Mowing | Triplets | 20.467 | 20.496 | 0.029 (-10.917, 12.812) | 6.131 | 0.005 | 0.996 |
| Grazing | Pairs | 31.826 | 36.170 | 4.344 (-18.901, 13.839) | 8.514 | 0.510 | 0.610 |
| Grazing | Triplets | 18.640 | 19.502 | 0.861 (-14.455, 9.241) | 6.196 | 0.139 | 0.889 |

**Table S11.** Endpoint comparisons of median structural niche differences (SND; A) and structural fitness differences (SFD; B) in Fig. S5. All three components use 1,135 transitions from 150 plots and a common 48-species pool (1,128 pairs and 17,296 triplets). Low and high refer to the endpoints in Table S10. Fitted changes compare endpoint medians; bias-corrected changes subtract estimated bootstrap bias. Bootstrap SE uses 999 successful joint parametric draws per component, refitting the population models described in Table S10. Calibration uses 750 datasets per component, with 199 inner refits attempted in each. Following a coverage diagnostic, all 12 intervals use a one-sided tolerance bound (Wilks, 1941) on the 95th percentile of calibration scores: the 728th ordered score, allowing for Monte Carlo uncertainty across the family. Intervals are the bias-corrected change ± this critical value times SE; $T$ is the absolute bias-corrected change divided by SE. Approximate two-sided $p$ values use corresponding exact-binomial upper bounds on simulated tail probabilities (Clopper & Pearson, 1934), with Holm adjustment across 12 comparisons. The 250 reserved validation datasets per component remain diagnostic checks, rather than fresh independent validation of this refinement. Endpoint comparisons do not test curve shape or isolate independent effects of correlated management practices.

A) Structural niche differences (SND)

| **Component** | **Assemblage** | **Fitted change** | **Bias-corrected change** | **Bootstrap SE** | ***T*** | **95% interval** | ***p*** |
|---|---|---|---|---|---|---|---|
| Fertilisation | Pairs | 0.017 | 0.047 | 0.042 | 1.128 | -0.071 to 0.165 | 1.000 |
| Fertilisation | Triplets | -0.125 | 0.068 | 0.083 | 0.818 | -0.363 to 0.499 | 1.000 |
| Mowing | Pairs | -0.045 | -0.032 | 0.036 | 0.895 | -0.128 to 0.063 | 1.000 |
| Mowing | Triplets | -0.281 | -0.212 | 0.069 | 3.091 | -0.446 to 0.021 | 0.983 |
| Grazing | Pairs | -0.010 | 0.005 | 0.033 | 0.147 | -0.090 to 0.099 | 1.000 |
| Grazing | Triplets | -0.250 | -0.190 | 0.067 | 2.817 | -0.423 to 0.043 | 1.000 |

B) Structural fitness differences (SFD)

| **Component** | **Assemblage** | **Fitted change** | **Bias-corrected change** | **Bootstrap SE** | ***T*** | **95% interval** | ***p*** |
|---|---|---|---|---|---|---|---|
| Fertilisation | Pairs | 19.176 | 37.378 | 15.335 | 2.437 | -20.959 to 95.715 | 1.000 |
| Fertilisation | Triplets | 8.391 | 23.691 | 10.571 | 2.241 | -18.479 to 65.860 | 1.000 |
| Mowing | Pairs | -4.110 | 0.299 | 14.883 | 0.020 | -50.100 to 50.698 | 1.000 |
| Mowing | Triplets | -9.435 | -5.520 | 10.257 | 0.538 | -39.064 to 28.023 | 1.000 |
| Grazing | Pairs | -1.916 | -6.225 | 16.280 | 0.382 | -68.401 to 55.951 | 1.000 |
| Grazing | Triplets | 2.189 | 6.217 | 11.554 | 0.538 | -32.596 to 45.030 | 1.000 |

**Table S12.** Management-component contrasts in community-pair differential and community-pair overlap corresponding to Fig. S6. Sections A, B and C show fertilisation, mowing and grazing, respectively; subsections 1 and 2 report community-pair differential and community-pair overlap. Low, middle and high denote fertilisation 0, 172.675 and 345.35 kg N $ha^{-1}$ $year^{-1}$; mowing 0, 2 and 4 cuts $year^{-1}$; and grazing 0, 822.084 and 1,644.167 livestock-unit days $ha^{-1}$ $year^{-1}$. Each contrast subtracts the earlier-level mean from the later-level mean. Estimates average paired contrasts from ten independent samples of 1,000 assemblages per size, using the common 48-species pool in Fig. S4. The same combinations are evaluated across levels and components. Monte Carlo standard errors (MC SE) and pointwise 95% Monte Carlo intervals follow Table S8. Agreement counts samples with the same sign as the mean contrast. These quantify combination-sampling precision, not population-model uncertainty. Positive differential changes indicate a shift towards greater promotion of feasibility; negative overlap changes indicate greater dependence on indirect effects. Overlap means exclude empty full feasibility domains separately at each level. Components are fitted separately, so these contrasts do not isolate independent management effects.

A) Fertilisation

*A.1) Community-pair differential*

| Species | Contrast | Change (95% MC interval) | MC SE | Agreement |
|---|---|---|---|---|
| 3 | Middle - low | 0.0042 (-0.0009, 0.0094) | 0.0023 | 8/10 |
| 3 | High - middle | 0.0171 (0.0133, 0.0210) | 0.0017 | 10/10 |
| 3 | High - low | 0.0214 (0.0168, 0.0260) | 0.0020 | 10/10 |
| 5 | Middle - low | -0.0180 (-0.0232, -0.0128) | 0.0023 | 10/10 |
| 5 | High - middle | 0.0238 (0.0196, 0.0281) | 0.0019 | 10/10 |
| 5 | High - low | 0.0058 (0.0001, 0.0116) | 0.0026 | 7/10 |
| 7 | Middle - low | -0.0216 (-0.0272, -0.0159) | 0.0025 | 10/10 |
| 7 | High - middle | 0.0147 (0.0118, 0.0175) | 0.0013 | 10/10 |
| 7 | High - low | -0.0069 (-0.0120, -0.0018) | 0.0023 | 8/10 |
| 11 | Middle - low | 0.0071 (0.0033, 0.0110) | 0.0017 | 9/10 |
| 11 | High - middle | 0.0035 (0.0028, 0.0043) | 0.0003 | 10/10 |
| 11 | High - low | 0.0107 (0.0072, 0.0142) | 0.0015 | 10/10 |

*A.2) Community-pair overlap*

| Species | Contrast | Change (95% MC interval) | MC SE | Agreement |
|---|---|---|---|---|
| 3 | Middle - low | -0.1603 (-0.1683, -0.1524) | 0.0035 | 10/10 |
| 3 | High - middle | -0.1676 (-0.1748, -0.1604) | 0.0032 | 10/10 |
| 3 | High - low | -0.3280 (-0.3366, -0.3193) | 0.0038 | 10/10 |
| 5 | Middle - low | -0.4024 (-0.4116, -0.3932) | 0.0041 | 10/10 |
| 5 | High - middle | -0.2509 (-0.2588, -0.2430) | 0.0035 | 10/10 |
| 5 | High - low | -0.6533 (-0.6599, -0.6467) | 0.0029 | 10/10 |
| 7 | Middle - low | -0.4990 (-0.5026, -0.4954) | 0.0016 | 10/10 |
| 7 | High - middle | -0.1343 (-0.1377, -0.1308) | 0.0015 | 10/10 |
| 7 | High - low | -0.6333 (-0.6359, -0.6307) | 0.0011 | 10/10 |
| 11 | Middle - low | -0.3248 (-0.3276, -0.3220) | 0.0012 | 10/10 |
| 11 | High - middle | -0.0101 (-0.0111, -0.0092) | 0.0004 | 10/10 |
| 11 | High - low | -0.3349 (-0.3376, -0.3323) | 0.0012 | 10/10 |

**Table S12 (continued).**

B) Mowing

*B.1) Community-pair differential*

| Species | Contrast | Change (95% MC interval) | MC SE | Agreement |
|---|---|---|---|---|
| 3 | Middle - low | -0.0112 (-0.0160, -0.0064) | 0.0021 | 10/10 |
| 3 | High - middle | 0.0293 (0.0261, 0.0326) | 0.0014 | 10/10 |
| 3 | High - low | 0.0181 (0.0141, 0.0222) | 0.0018 | 10/10 |
| 5 | Middle - low | -0.0507 (-0.0535, -0.0479) | 0.0012 | 10/10 |
| 5 | High - middle | 0.0381 (0.0346, 0.0416) | 0.0016 | 10/10 |
| 5 | High - low | -0.0126 (-0.0156, -0.0096) | 0.0013 | 10/10 |
| 7 | Middle - low | -0.0628 (-0.0686, -0.0571) | 0.0026 | 10/10 |
| 7 | High - middle | 0.0379 (0.0350, 0.0408) | 0.0013 | 10/10 |
| 7 | High - low | -0.0250 (-0.0295, -0.0205) | 0.0020 | 10/10 |
| 11 | Middle - low | -0.0220 (-0.0251, -0.0189) | 0.0014 | 10/10 |
| 11 | High - middle | 0.0177 (0.0166, 0.0187) | 0.0005 | 10/10 |
| 11 | High - low | -0.0043 (-0.0072, -0.0014) | 0.0013 | 9/10 |

*B.2) Community-pair overlap*

| Species | Contrast | Change (95% MC interval) | MC SE | Agreement |
|---|---|---|---|---|
| 3 | Middle - low | -0.0685 (-0.0738, -0.0631) | 0.0024 | 10/10 |
| 3 | High - middle | -0.2238 (-0.2277, -0.2200) | 0.0017 | 10/10 |
| 3 | High - low | -0.2923 (-0.2984, -0.2861) | 0.0027 | 10/10 |
| 5 | Middle - low | -0.1443 (-0.1547, -0.1338) | 0.0046 | 10/10 |
| 5 | High - middle | -0.4045 (-0.4149, -0.3941) | 0.0046 | 10/10 |
| 5 | High - low | -0.5487 (-0.5605, -0.5370) | 0.0052 | 10/10 |
| 7 | Middle - low | -0.1972 (-0.2046, -0.1899) | 0.0032 | 10/10 |
| 7 | High - middle | -0.3522 (-0.3597, -0.3448) | 0.0033 | 10/10 |
| 7 | High - low | -0.5495 (-0.5524, -0.5465) | 0.0013 | 10/10 |
| 11 | Middle - low | -0.1658 (-0.1706, -0.1609) | 0.0021 | 10/10 |
| 11 | High - middle | -0.0932 (-0.0962, -0.0902) | 0.0013 | 10/10 |
| 11 | High - low | -0.2589 (-0.2619, -0.2559) | 0.0013 | 10/10 |

**Table S12 (continued).**

C) Grazing

*C.1) Community-pair differential*

| Species | Contrast | Change (95% MC interval) | MC SE | Agreement |
|---|---|---|---|---|
| 3 | Middle - low | 0.0221 (0.0180, 0.0262) | 0.0018 | 10/10 |
| 3 | High - middle | 0.0119 (0.0088, 0.0150) | 0.0014 | 10/10 |
| 3 | High - low | 0.0340 (0.0293, 0.0387) | 0.0021 | 10/10 |
| 5 | Middle - low | 0.0265 (0.0231, 0.0300) | 0.0015 | 10/10 |
| 5 | High - middle | -0.0070 (-0.0093, -0.0046) | 0.0010 | 10/10 |
| 5 | High - low | 0.0195 (0.0157, 0.0234) | 0.0017 | 10/10 |
| 7 | Middle - low | 0.0161 (0.0126, 0.0195) | 0.0015 | 10/10 |
| 7 | High - middle | -0.0083 (-0.0110, -0.0057) | 0.0012 | 10/10 |
| 7 | High - low | 0.0077 (0.0052, 0.0103) | 0.0011 | 10/10 |
| 11 | Middle - low | 0.0156 (0.0131, 0.0182) | 0.0011 | 10/10 |
| 11 | High - middle | -0.0006 (-0.0010, -0.0001) | 0.0002 | 8/10 |
| 11 | High - low | 0.0151 (0.0126, 0.0176) | 0.0011 | 10/10 |

*C.2) Community-pair overlap*

| Species | Contrast | Change (95% MC interval) | MC SE | Agreement |
|---|---|---|---|---|
| 3 | Middle - low | -0.1595 (-0.1654, -0.1536) | 0.0026 | 10/10 |
| 3 | High - middle | -0.1869 (-0.1930, -0.1808) | 0.0027 | 10/10 |
| 3 | High - low | -0.3464 (-0.3554, -0.3374) | 0.0040 | 10/10 |
| 5 | Middle - low | -0.3988 (-0.4054, -0.3921) | 0.0029 | 10/10 |
| 5 | High - middle | -0.2364 (-0.2410, -0.2318) | 0.0020 | 10/10 |
| 5 | High - low | -0.6352 (-0.6410, -0.6294) | 0.0026 | 10/10 |
| 7 | Middle - low | -0.4653 (-0.4696, -0.4609) | 0.0019 | 10/10 |
| 7 | High - middle | -0.1051 (-0.1102, -0.1000) | 0.0023 | 10/10 |
| 7 | High - low | -0.5704 (-0.5752, -0.5655) | 0.0021 | 10/10 |
| 11 | Middle - low | -0.2080 (-0.2101, -0.2059) | 0.0009 | 10/10 |
| 11 | High - middle | -0.0061 (-0.0064, -0.0058) | 0.0001 | 10/10 |
| 11 | High - low | -0.2141 (-0.2162, -0.2120) | 0.0009 | 10/10 |

**Table S13.** Differences in community-pair differential (A) and community-pair overlap (B) between adjacent assemblage sizes, corresponding to Fig. S6. Low, middle and high are the component levels defined in Table S12. Each size contrast subtracts the smaller-assemblage mean from the larger-assemblage mean (e.g., 5 minus 3 species). Estimates average contrasts from ten independent sampling runs, each containing 1,000 assemblages per size from the common 48-species pool in Fig. S4. Monte Carlo standard errors (MC SE) and pointwise 95% Monte Carlo intervals follow Table S8. Agreement counts runs with the same sign as the mean contrast. These quantify combination-sampling precision, not population-model uncertainty. Positive differential differences indicate a shift towards greater promotion of feasibility in larger assemblages; negative overlap differences indicate greater dependence on indirect effects. Overlap means exclude empty full feasibility domains separately at each size and level. Assemblages are sampled separately at each size, so these comparisons do not represent adding species to a particular assemblage. Components are fitted separately and do not isolate independent management effects.

A) Community-pair differential

| Component | Size contrast | Level | Difference (95% MC interval) | MC SE | Agreement |
|---|---|---|---|---|---|
| Fertilisation | 5 - 3 | Low | 0.0129 (0.0102, 0.0157) | 0.0012 | 10/10 |
| Fertilisation | 5 - 3 | Middle | -0.0093 (-0.0177, -0.0009) | 0.0037 | 8/10 |
| Fertilisation | 5 - 3 | High | -0.0026 (-0.0090, 0.0038) | 0.0028 | 5/10 |
| Fertilisation | 7 - 5 | Low | -0.0066 (-0.0128, -0.0003) | 0.0028 | 8/10 |
| Fertilisation | 7 - 5 | Middle | -0.0102 (-0.0147, -0.0056) | 0.0020 | 10/10 |
| Fertilisation | 7 - 5 | High | -0.0193 (-0.0235, -0.0152) | 0.0018 | 10/10 |
| Fertilisation | 11 - 7 | Low | -0.0252 (-0.0303, -0.0201) | 0.0023 | 10/10 |
| Fertilisation | 11 - 7 | Middle | 0.0035 (-0.0002, 0.0072) | 0.0016 | 8/10 |
| Fertilisation | 11 - 7 | High | -0.0076 (-0.0095, -0.0058) | 0.0008 | 10/10 |
| Mowing | 5 - 3 | Low | 0.0238 (0.0207, 0.0269) | 0.0014 | 10/10 |
| Mowing | 5 - 3 | Middle | -0.0156 (-0.0208, -0.0105) | 0.0023 | 10/10 |
| Mowing | 5 - 3 | High | -0.0069 (-0.0105, -0.0032) | 0.0016 | 9/10 |
| Mowing | 7 - 5 | Low | -0.0035 (-0.0092, 0.0021) | 0.0025 | 7/10 |
| Mowing | 7 - 5 | Middle | -0.0157 (-0.0196, -0.0118) | 0.0017 | 10/10 |
| Mowing | 7 - 5 | High | -0.0160 (-0.0184, -0.0135) | 0.0011 | 10/10 |
| Mowing | 11 - 7 | Low | -0.0274 (-0.0332, -0.0216) | 0.0026 | 10/10 |
| Mowing | 11 - 7 | Middle | 0.0134 (0.0105, 0.0163) | 0.0013 | 10/10 |
| Mowing | 11 - 7 | High | -0.0068 (-0.0080, -0.0055) | 0.0006 | 10/10 |
| Grazing | 5 - 3 | Low | 0.0093 (0.0067, 0.0120) | 0.0012 | 10/10 |
| Grazing | 5 - 3 | Middle | 0.0138 (0.0094, 0.0182) | 0.0019 | 10/10 |
| Grazing | 5 - 3 | High | -0.0051 (-0.0106, 0.0005) | 0.0024 | 7/10 |
| Grazing | 7 - 5 | Low | -0.0118 (-0.0158, -0.0077) | 0.0018 | 9/10 |
| Grazing | 7 - 5 | Middle | -0.0222 (-0.0268, -0.0177) | 0.0020 | 10/10 |
| Grazing | 7 - 5 | High | -0.0236 (-0.0266, -0.0206) | 0.0013 | 10/10 |
| Grazing | 11 - 7 | Low | -0.0200 (-0.0229, -0.0171) | 0.0013 | 10/10 |
| Grazing | 11 - 7 | Middle | -0.0204 (-0.0231, -0.0176) | 0.0012 | 10/10 |
| Grazing | 11 - 7 | High | -0.0126 (-0.0136, -0.0117) | 0.0004 | 10/10 |

**Table S13 (continued).**

B) Community-pair overlap

| Component | Size contrast | Level | Difference (95% MC interval) | MC SE | Agreement |
|---|---|---|---|---|---|
| Fertilisation | 5 - 3 | Low | -0.1287 (-0.1303, -0.1271) | 0.0007 | 10/10 |
| Fertilisation | 5 - 3 | Middle | -0.3708 (-0.3850, -0.3566) | 0.0063 | 10/10 |
| Fertilisation | 5 - 3 | High | -0.4540 (-0.4665, -0.4416) | 0.0055 | 10/10 |
| Fertilisation | 7 - 5 | Low | -0.1652 (-0.1679, -0.1625) | 0.0012 | 10/10 |
| Fertilisation | 7 - 5 | Middle | -0.2618 (-0.2715, -0.2522) | 0.0043 | 10/10 |
| Fertilisation | 7 - 5 | High | -0.1452 (-0.1512, -0.1393) | 0.0026 | 10/10 |
| Fertilisation | 11 - 7 | Low | -0.3338 (-0.3369, -0.3307) | 0.0014 | 10/10 |
| Fertilisation | 11 - 7 | Middle | -0.1596 (-0.1631, -0.1562) | 0.0015 | 10/10 |
| Fertilisation | 11 - 7 | High | -0.0355 (-0.0365, -0.0345) | 0.0005 | 10/10 |
| Mowing | 5 - 3 | Low | -0.1609 (-0.1645, -0.1574) | 0.0016 | 10/10 |
| Mowing | 5 - 3 | Middle | -0.2367 (-0.2493, -0.2241) | 0.0056 | 10/10 |
| Mowing | 5 - 3 | High | -0.4174 (-0.4308, -0.4040) | 0.0059 | 10/10 |
| Mowing | 7 - 5 | Low | -0.1899 (-0.1925, -0.1872) | 0.0012 | 10/10 |
| Mowing | 7 - 5 | Middle | -0.2429 (-0.2567, -0.2290) | 0.0061 | 10/10 |
| Mowing | 7 - 5 | High | -0.1906 (-0.2016, -0.1796) | 0.0049 | 10/10 |
| Mowing | 11 - 7 | Low | -0.3429 (-0.3478, -0.3381) | 0.0021 | 10/10 |
| Mowing | 11 - 7 | Middle | -0.3115 (-0.3187, -0.3042) | 0.0032 | 10/10 |
| Mowing | 11 - 7 | High | -0.0524 (-0.0548, -0.0500) | 0.0010 | 10/10 |
| Grazing | 5 - 3 | Low | -0.1596 (-0.1626, -0.1566) | 0.0013 | 10/10 |
| Grazing | 5 - 3 | Middle | -0.3988 (-0.4067, -0.3909) | 0.0035 | 10/10 |
| Grazing | 5 - 3 | High | -0.4483 (-0.4600, -0.4366) | 0.0052 | 10/10 |
| Grazing | 7 - 5 | Low | -0.2015 (-0.2050, -0.1979) | 0.0016 | 10/10 |
| Grazing | 7 - 5 | Middle | -0.2680 (-0.2751, -0.2608) | 0.0032 | 10/10 |
| Grazing | 7 - 5 | High | -0.1367 (-0.1423, -0.1310) | 0.0025 | 10/10 |
| Grazing | 11 - 7 | Low | -0.3811 (-0.3852, -0.3769) | 0.0018 | 10/10 |
| Grazing | 11 - 7 | Middle | -0.1238 (-0.1276, -0.1201) | 0.0017 | 10/10 |
| Grazing | 11 - 7 | High | -0.0248 (-0.0274, -0.0221) | 0.0012 | 10/10 |

#### Software used

Analyses used R 4.3.2 on the Hyperion cluster at the Donostia International Physics Center (DIPC) and R 4.4.3 locally (R Core Team, 2025). Packages used for the analyses, figure preparation and software citation collection were `cowplot` v. 1.2.0 (Wilke, 2025); `emmeans` v. 1.10.1 (Lenth, 2024); `ggh4x` v. 0.3.0 (van den Brand, 2024); `ggplot2` v. 4.0.3 (Wickham, 2016); `grateful` v. 0.3.0 (Rodríguez-Sánchez & Jackson, 2025); `gridExtra` v. 2.3 (Auguie, 2017); `gtable` v. 0.3.6 (Wickham & Pedersen, 2024); `lpSolve` versions 5.6.19 and 5.6.21 (Berkelaar, 2024); `nlme` versions 3.1-164 and 3.1-167 (Pinheiro et al., 2023, 2025); `patchwork` v. 1.3.2 (Pedersen, 2025); `plyr` v. 1.8.9 (Wickham, 2011); `ragg` v. 1.3.1 (Pedersen & Shemanarev, 2024); `Rcpp` versions 1.0.11 and 1.1.1-1.1 (Eddelbuettel & François, 2011); `scales` v. 1.4.0 (Wickham et al., 2025); `svglite` v. 2.1.3 (Wickham et al., 2023). Documents were produced using Quarto 1.8.27 (Allaire et al., 2026) and Typst 0.13.0 (Mädje et al., 2025).
